# Leveraging High-dimensional Seed Colorspace in Phenomic Selection Models for Herbaceous Perennial Crops

**DOI:** 10.1101/2025.07.28.667321

**Authors:** William Keaggy, Zachary N Harris, Jackson Braley, Eric Cassetta, Jorge Gutierrez, Emelyn Piotter, Haley Schuhl, Jared L Crain, Lee R DeHaan, M. Kathryn Turner, Noah Fahlgren, Jenna Hershberger, Brandon Schlautman, David L Van Tassel, Matthew J Rubin, Allison J Miller

**Author notes:** These authors contributed equally to this work (William Keaggy, Zachary N Harris).

## Abstract

Herbaceous perennial crops, which only recently entered the domestication pipeline, offer the potential of ecological benefits in agricultural systems but remain underdeveloped, partly due to slow breeding cycles. Early-stage phenomic selection, the process of using high-dimensional secondary traits to predict target traits, may accelerate improvement. We evaluated whether seed traits derived from image scans could be used to predict germination traits in three perennial crop candidate species: sainfoin (*Onobrychis viciifolia*), intermediate wheatgrass (*Thinopyrum intermedium*), and silflower (*Silphium integrifolium*). Over 20,000 seeds were scanned, and PlantCV was used to extract 692 hue, saturation, and value (HSV) color features along with five morphological traits. We demonstrated that seed color, morphology, and germination traits were all influenced by maternal family, but patterns of correlation between germination traits and color and morphology traits are complex. To enable prediction from high-dimensional seed traits, we constructed relationship matrices from combinations of HSV and morphology features for use in phenomic selection models to predict germination proportion and timing. Model performance varied by trait, species, and cohort, and in many cases, predictions were significantly better than chance. HSV features, when combined with morphology, often yielded the most accurate predictions, with performance reaching a maximum of r = 0.33. These results demonstrate the potential of low-cost, image-based phenomic data to inform early-stage selection and support the development of sustainable perennial crops.

**Plain language summary:** Many of the world’s most important crops (e.g., wheat, maize, soy) live for less than one year, leaving soil bare for many months and subject to erosion. Longer-lived (perennial) alternatives to these crops have the potential to support a more sustainable agricultural system by improving soil health and reducing the need for chemical inputs. However, perennial herbaceous plants were not domesticated by early farmers, and now lag behind annual crops in development. They take longer to breed and often lack the genetic resources needed for modern breeding approaches. In this study, we tested whether simple seed images derived from color scanners could be used to predict key traits like germination proportion and timing in emerging perennial crops. We found that seed color and shape varied by maternal family and that color, especially when combined with seed size and shape, could help predict germination performance. These predictions were significantly better than chance. This work shows that low-cost, image-based seed data can support early-stage selection in breeding programs, especially for undeveloped perennial crops, helping save time, reduce cost, and accelerate progress toward more sustainable agriculture.

## Introduction

Although most of the plant world is composed of species that live for multiple years (perennial species), agriculture is dependent primarily on plants that live for one year (annual species), including nearly all grains and legumes (Friedman, 2020; Friedman & Rubin, 2015; Kreitzman et al., 2020; Pimentel et al., 2012; Poppenwimer et al., 2023). Annual crops grown in monoculture, which must be replanted every year and typically leave soil bare for part of the year, contribute to erosion and accelerated soil degradation (Boulal et al., 2011; Thaler et al., 2022). While woody perennials (fruit and nut trees) are cultivated at scale, herbaceous perennials have not been widely domesticated (Ciotir et al., 2019; Van Tassel et al., 2010). Herbaceous perennial grains offer a unique opportunity to develop agricultural systems that provide economically viable products and important ecosystem services, as the deep and persistent multiyear root systems of herbaceous perennials increase soil fertility (Freibauer et al., 2004) and reduce erosion and runoff (Ditsch, 1998). Perennial herbaceous grains also have the potential to reduce the cost of agriculture through less chemical input and lower labor requirements (Crews et al., 2018; Zhang et al., 2022).

The long life cycles and complex life histories of many perennial herbaceous species slow the breeding process relative to annuals, contributing to the historical emphasis on annual crops (Schaart et al., 2016; Schlautman et al., 2018). For example, perennials may initially favor resource allocation to vegetative growth, laying the foundation for greater future flowering opportunities in subsequent seasons (Friedman & Rubin, 2015). Yearly seed production in perennial species often negatively correlates with lifespan, where longer-lived species produce fewer seeds per year than annuals (Obeso, 2002), and some species exhibit high interannual variability in seed output. These biological constraints likely made perennials less attractive to early agriculturalists, contributing to a developmental lag of up to 10,000 years relative to major annual crops. These challenges, coupled with the growing ecological and economic demand for sustainable cropping systems, underscore the need for innovative breeding techniques to support perennial grain development.

In major annual crops such as maize and soybean, genomic selection (GS) enables predictive breeding using genome-wide markers (Lorenz et al., 2011; Meuwissen et al., 2001; Voss-Fels et al., 2019). During GS, a training population with paired phenotypic data and genome-wide markers is used to develop a model that maps marker variation to estimated phenotypes or breeding values, often through the construction of relationship matrices or richly parameterized linear mixed models (Heffner et al., 2009; Van Tassel et al., 2022). Phenotypes for new individuals that have genome-wide marker data but not phenotypic data can be estimated using those trained models. Unfortunately, GS can be impractical for emerging perennial crops, which often have large, heterozygous, and frequently polyploid genomes as well as obligate outcrossing mating systems (Dudash & Fenster, 2001; Stevens et al., 2020; Veeckman et al., 2019). Despite concerns about the pace of domesticating new crops, recent efforts in perennial rice, intermediate wheatgrass, and silflower show that selection for traits like yield is likely achievable over relatively short timeframes when resources are available (Crain et al., 2021; DeHaan et al., 2020; Schaart et al., 2016; Van Tassel et al., 2017; Zhang et al., 2022).

Phenomic selection (PS) is an emerging tool with potential to accelerate breeding in species where genomic resources are either non-existent or underdeveloped. Unlike GS, which relies on genetic markers, PS replaces genome-wide marker data with high-dimensional phenotypic data (often called secondary traits, such as hyperspectral reflectance) to predict traits of interest (often called primary traits, for example, yield) (Rincent et al., 2018). This approach has been successfully used to predict germination, seedling vigor, and mature plant performance in both annual and perennial systems (Krause et al., 2019; Laurençon et al., 2024; Robert, Goudemand, et al., 2022; Tu et al., 2024; Van Tassel et al., 2022). In a study of winter wheat, PS outperformed GS in nearly half of the yield predictions and 62% of the heading date predictions (Robert, Auzanneau, et al., 2022). In both this study and another on bread wheat, results were strongest when both methods, PS and GS, were combined (Krause et al., 2019; Robert, Brault, et al., 2022). In maize, PS has performed comparably to GS, while being more cost-effective and less sensitive to population structure (Weiß et al., 2022). While PS has shown promise in major grain crops, open questions remain about what secondary traits can be used to predict relevant primary traits, the optimal source of secondary traits from which to make predictions, and whether extensive variation in secondary traits influences the ability to make phenomic predictions.

Ongoing research continues to explore different sources of high-dimensional phenomic data suitable for phenomic selection models (Van Tassel et al., 2022). For example, image-based color data, including seed color, has been widely used in agricultural research as it reflects the underlying physiology and biochemistry of the plant (Gebregziabher et al., 2022; Nayak et al., 2020). One promising format for color data is the HSV (hue, saturation, value) colorspace, where hue represents the color itself (e.g., red, yellow, green), saturation describes the intensity or vibrancy of the color, and value reflects its brightness or lightness (Shuhua & Gaizhi, 2010). These three components define a multidimensional colorspace from which relationships can be estimated and used in PS.

In this study, we use HSV color profiles and seed morphological traits extracted from images to predict germination traits, with the broader goal of improving phenomic selection pipelines for emerging perennial crops. We focus on three candidate species being developed as perennial grains or oilseeds: *Onobrychis viciifolia* Scop, *Thinopyrum intermedium* (Host) Barkworth & D.R.Dewey, and *Silphium integrifolium* Michx, which span a range of seed shape, color, and morphology. *Onobrychis viciifolia* (Baki™ bean, hereafter sainfoin) is a leguminous species native to eastern and central Asia (Hayot Carbonero et al., 2011). Sainfoin has been widely used as a forage crop across Europe (Frame et al., 1998; Hayot Carbonero et al., 2011), but has recently gained renewed interest for human consumption, as it is noted for its dietary and environmental benefits (Craine et al., 2023; Schlautman et al., 2018). *Thinopyrum intermedium* (Kernza**®**/intermediate wheatgrass, hereafter IWG) is a relative of wheat that is native to the Mediterranean regions of Europe and Asia (Wagoner, 1990). Originally used as a forage crop, IWG has been under selection for improved grain yield as a target for perennial grain similar to wheat since the 1980s (Bajgain et al., 2022; Smaje, 2015). *Silphium integrifolium* (hereafter silflower) is a sunflower relative native to central North America. Silflower is being domesticated as a dual-use oilseed and forage crop (Vilela et al., 2020).

Using sainfoin, IWG, and silflower as model systems, we investigate the efficacy of HSV color profiles in phenomic selection. Specifically, we address three questions: (1) Within each species, how variable are seed color and morphology within and across populations? (2) Do seed color and morphology correlate with germination traits? (3) Can these traits generate high-dimensional relationship matrices that accurately predict germination proportion and germination timing? This work presents the first comprehensive analysis of high-dimensional seed color data in emerging perennial crops and evaluates its utility for PS.

## Methods

### Experimental design

To assess variation in seed traits and the capacity of seed traits to predict germination, this experiment included two to four experimental cohorts for each of three perennial herbaceous species: sainfoin, IWG, and silflower. Each study species consisted of multiple cohorts, with seeds in each cohort sourced from different breeding cycles and/or half-sib maternal families. We acquired seeds of each species from their respective breeding programs at The Land Institute in Salina, KS. IWG is discussed in terms of breeding cycles which have been documented by (Bajgain et al., 2022), but both sainfoin and silflower are comparatively early in their respective breeding programs.

Sainfoin was evaluated in two cohorts, containing 60 and 64 half-sib maternal families, respectively, with 14 to 42 seeds per family (n_1_ = 2520, n_2_ = 2582, Table 1). Each cohort is derived from a set of shared accessions, but several rounds of outcrossing have resulted in a less structured population. We planted the first cohort in the summer of 2021 and the second in the summer of 2022. IWG was evaluated in two cohorts containing 95 and 96 half-sib maternal families, respectively, with 26 seeds per family (n_1_ = 2470, n_2_ = 2496; Table 1). Seeds from each cohort were acquired from the TLI breeding cycles 11 and 12, where TLI-Cycle 12 are progeny from selected, intermated TLI-Cycle 11 plants. We planted the first cohort in the summer of 2021 and the second in the summer of 2022. A total of 2,280 IWG individuals from cohorts 1 and 2 were genotyped using genotyping-by-sequencing (GBS) following the protocols of (Poland et al., 2012). Bioinformatic processing followed the same procedure that the TLI IWG breeding program has used historically (Crain et al., 2021), yielding approximately 15,000 single-nucleotide polymorphisms (SNPs) per sample. Using the genotyped individuals, we were able to assign paternity to complete full pedigrees of the 2,280 IWG individuals using Cervus software (Kalinowski et al., 2007), obtaining both maternal and paternal families. In previous work in IWG, this method of paternity analysis has shown over 90% agreement with written records in controlled crosses (Crain et al., 2020). Silflower was assessed in four cohorts, representing 31, 62, 147, and 44 half-sib maternal families, respectively (n_1_ = 3835, n_2_ = 2169, n_3_ = 2192, n_4_ = 2376). Cohort 2 was derived from controlled crosses of cohort 1. Cohort 3 was a set of open-pollinated crosses from cohort 1. Cohort 4 was derived from a diverse set of crosses unrelated to cohort 1. We planted the first cohort in the spring of 2022 and cohorts 2, 3, and 4 in the summer of 2024.

**Table 1.**
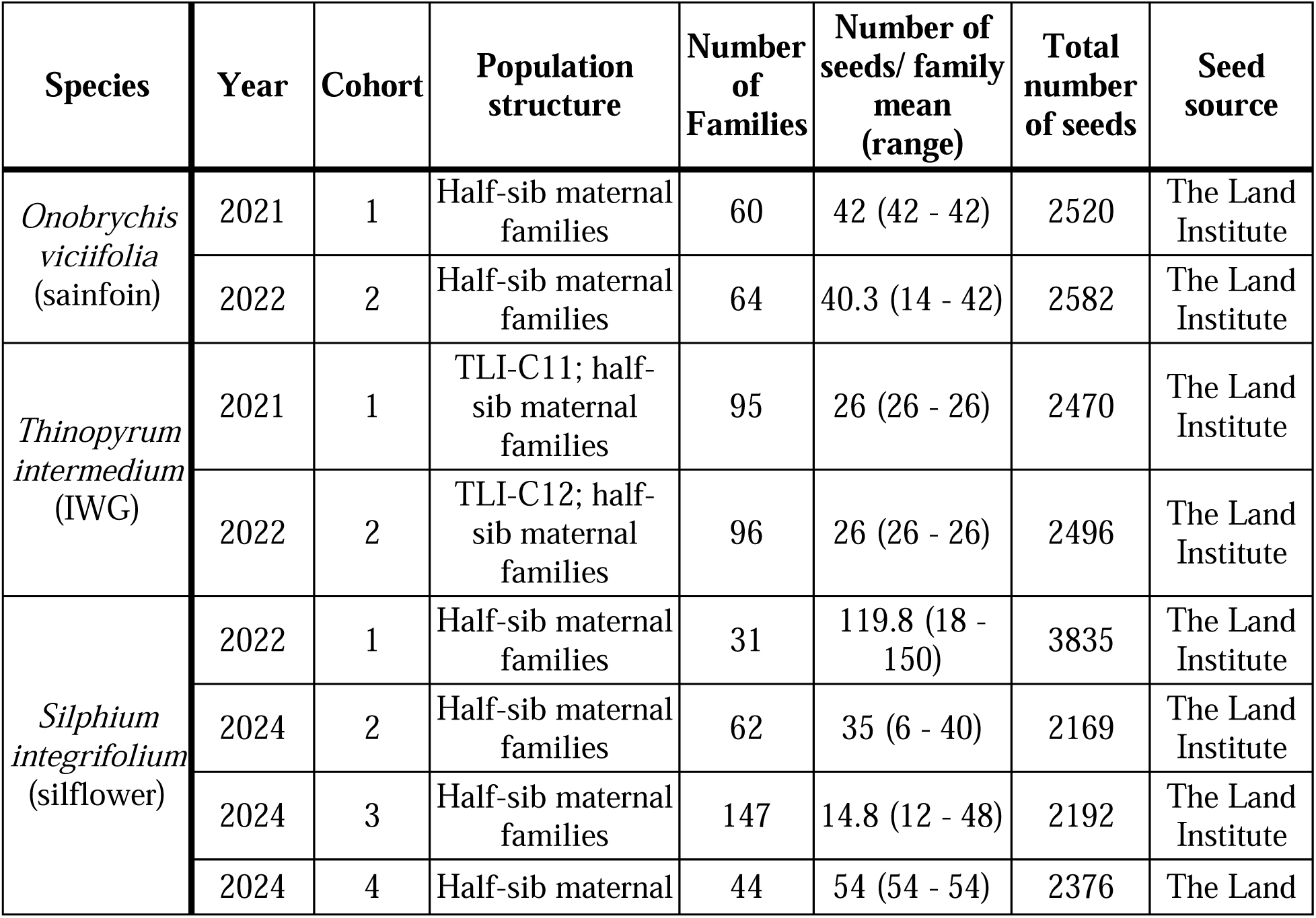
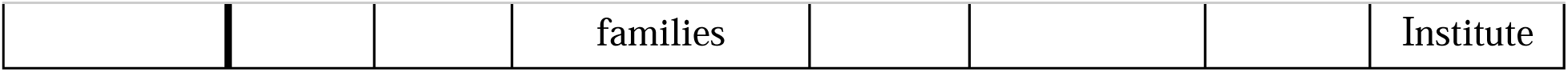
Cohort information for three study species.

### Seed Data

In this paper, we refer to the tissues of sainfoin and silflower from which we gathered data as seeds, although we acknowledge the proper term is fruit, as in both species the seeds are enclosed in maternal tissue. To generate high-dimensional color and morphology traits for seeds, each seed was assigned a unique ID, placed on a grid layout, and scanned using an EPSON DS-50000 scanner (Nagano, Japan) at 1200 DPI. A SpyderCheckr® 24 color card (datacolor, Lawrenceville, New Jersey, United States) and a ruler were included in each TIFF image. Silflower seeds were placed ventral side down due to the asymmetry of the seed. Example seed images can be found in Supplemental Figure 1. After scanning, each seed was placed in a microcentrifuge tube or 24-well plate labeled with the seed ID.

We analyzed each seed image using PlantCV (Version 4.0 for sainfoin and IWG, version 4.5.1 for silflower), a Python-based, open-source image analysis software package (Fahlgren et al., 2015; Gehan et al., 2017). We evaluated each channel in HSV and LAB (channels: L = lightness, A = red/green value, B = blue/yellow value) colorspaces to segment seeds from the white background. Saturation almost perfectly isolated each seed from the white background across all three species.

Each scan of sainfoin and IWG was cropped to include only the grid of seeds. Silflower scans were cropped into individual seed scans. These scans were converted to binary images using the plantcv.threshold.otsu function and cleaned using the fill function to remove any objects in the image less than 500 pixels (∼0.22 mm^2^) for IWG and 5000 pixels (∼2.2 mm^2^) for sainfoin and silflower. The clean binary image was applied as a mask to the original color image to isolate seeds from the background. We created a grid of regions of interest (ROI) using plantcv.roi.auto_grid function to align with the grid we created on the flatbed scanner. We used plantcv.create_labels to label seeds as objects within the image, and each labeled seed object was assigned to the corresponding ROI using plantcv.roi.filter. Seed morphological traits (area, length, width, perimeter, and circularity of each seed) were obtained from scans of each species using the plantcv.analyze.size function. Each morphological trait was measured in pixels, except for circularity, which was measured on a scale of 0 to 1 (0 = circle, 1 = ellipse). For seed color data, seed scans were used to estimate hue, saturation, and value (HSV) using the plantcv.analyze_color function. Color data included 692 HSV color traits: 180 hue features measured in 2-degree increments (0–360°) and 256 saturation and value features each, scaled from 0 to 1 based on 8-bit image values (0 - 255) (Agoston, 2005). As silflower used an updated version of PlantCV, the same features were extracted using the plantcv.analyze.size and plantcv.analyze.color functions to gather morphology and color data. All PlantCV workflows were developed in Jupyter Notebooks (Kluyver et al., 2016).

### Pre-Planting Treatments

Following scanning, seeds were germinated according to best practices for that species. Sainfoin seeds were not treated between scanning and planting. IWG seeds were soaked in water for 24 hours at room temperature. To remove the water, we placed small pieces of cotton in the top section of each tube. When we flipped the tubes, the water was absorbed into the cotton. We left the cotton in the tubes to maintain a humid environment until it was time to plant. We then wrapped tubes in foil to block out any light and cold treated them at 4°C for eight days before planting. Silflower seeds were treated with a chemical soak of 1 mM ethephon solution (4 mL of Florel (Montere®, Fresno, CA, Batch LG-030320A) to 1 L of DI water). We added enough solution to submerge each seed and allowed the seeds to soak for 24 hours at room temperature. For silflower cohort 1, we removed the solution using the previously described cotton method. For silflower cohorts 2-4, we placed a sieve on top of the open plate and decanted the solution. A trace amount of solution remained to maintain humidity. We then covered the trays in foil to block the light and cold treated them at 4°C for 12 days for cohort 1, and 13 days for cohorts 2-4.

### Planting

After pre-planting treatments, we planted seeds into square 7.6 cm wide by 20.3 cm tall MT38 Steuwe and Sons pots with capmat insert, filled with Berger Bm7 35% Bark HP + 14-14-14 Osmocote (1.5 lb/cu yd) soil (Saint-Modeste, Quebec, Canada). We planted two seeds per pot, placing seeds just under the soil surface. Pots were labeled with two barcodes that represented seed ID and transferred to a Conviron MTPC144 growth chamber (Winnipeg, Manitoba, Canada). Inside the chamber, pots were bottom watered daily and gently misted with DI water, twice daily, to saturation. The photoperiod was 13 hours, with 300 lumens of light, 60% humidity, a day temperature of 25°C, and a night temperature of 20°C.

### Germination

In this study, the emergence of any part of the seedling from the seed, and our observation of that seedling at or above the soil surface, was considered to be germination. We recorded germination daily, noting the number of days after planting that each seed germinated. The germination window varied across species, ranging from 10 to 15 days (sainfoin), 9 to 12 days (IWG), and 16 to 22 days (silflower). We calculated the germination proportion for each maternal family, as well as the time to germination for each seed.

### Data Analysis

To quantify existing variation in seed color and morphology within and across populations of the three focal species, we conducted statistical analyses in R (R version 4.4.2, 2024-10-31). Outliers in morphology and days to germination that were 1.5 times the interquartile range (IQR) above the third quartile or below the first quartile were removed. Seed HSV features were filtered to remove those with zero variance or those with only two unique values across all samples, as these likely reflected imaging artifacts or background noise rather than meaningful biological variation. For each continuous trait (area, width, length, perimeter, circularity, median hue, days to germination, and each frequency of hue, saturation, and value), we fit linear models parameterized with maternal parent ID to test for variation from maternal family on scaled and centered phenotypes. In the case of germination proportion, a binomial trait, we used a generalized linear model with a binomial link function and reported the chi-squared value. In the case of known paternity in IWG from genotypic data, we additionally fit linear models parameterized with paternal family as a single fixed effect. We then used the ggpairs function from GGally (v2.2.1; (Schloerke et al., 2024)) to plot family mean distributions of morphology traits, germination proportion, and days to germination and scatterplots showing the relationships between pairs of these traits. Pearson’s correlation coefficients were calculated to quantify the strength of the relationship for each pair of traits, separated by cohort and with cohorts combined.

To further characterize variation in seed color, we tested the extent to which each maternal lineage (and paternal family in the case of IWG) could be predicted from seed HSV data. We used gradient-boosted classification trees implemented in lightGBM (v4.5.0; (Shi et al., 2025)) to assign individual seeds to their maternal family based on filtered HSV features in a 5-fold cross-validation framework. We reported results as a measure of model accuracy relative to a null model, which only predicted the most frequent maternal family in each fold. We conducted one-tailed t-tests to determine whether a model’s predictive ability was significantly higher than that of the null model.

To test the capacity of seed characteristics to predict germination traits, we used a high-dimensional HSV color dataset grouped as hue, saturation, and value alone (H, S, or V), pairwise combinations of the three (HS, HV, SV), and all three together (HSV), along with seed morphology traits alone (Morph) and HSV combined with seed morphology (HSVMorph) to generate relationship matrices based on cosine similarity for all individuals in each species cohort. Following the methods of Krause et al. (2019), we used the BGLR package in R (v1.1.3; (Pérez & de los Campos, 2014)) to create a 5-fold cross-validation scheme and predict the germination proportion and time to germinate for each seed in the experiment. We used Pearson’s correlation to compare our predicted and observed values as a measure of model performance. We conducted a one-tailed t-test to determine if correlations were significantly larger than 0. Following the model fit, performance was assessed within each cohort using a linear model to determine whether different relationship matrices yielded different performances.

## Results

### Variation in seed HSV and morphology

We evaluated whether seed color and morphology traits captured genetic structure across three perennial crop species by analyzing how these traits vary among maternal families in multiple experimental cohorts. Across species, HSV color features consistently reflected strong maternal family differentiation. Morphological traits also varied significantly, though patterns differed across cohorts. Machine learning models using HSV data effectively predicted maternal family, with the highest accuracies in IWG and sainfoin. Silflower exhibited lower predictive performance but still showed substantial family-level structuring. These results demonstrate that high-dimensional seed traits encode detectable genetic signals and hold potential for early-stage PS. Below, we detail these findings by species and cohorts.

Sainfoin maternal family explained substantial variation in HSV features (Figure 1, Supplemental Figure 2). In hue features, the strongest maternal family differentiation was identified in features <50°, with cohort 1 showing additional describable variation around 250°, but cohort 2 only differentiating after 300°. In saturation features, cohort 1 exhibited a peak in family stratification around saturation = 0.55. Cohort 2 showed a similar, though broader, peak in stratification around saturation = 0.47. Overall, variation from maternal family was strongest in features relating to value, especially in cohort 2, peaking near value = 0.66. Using all HSV features, LightGBM models predicted maternal families with 84.9% and 96.5% accuracy in cohorts 1 and 2, respectively (Figure 2). Important features from the LightGBM aligned well with linear model peaks (Supplemental Figure 3). The optimal models for predicting maternal family in both cohorts outperformed null expectations (p < 0.001). We also found that seed morphology traits varied by maternal family (Figure 3; Supplemental Figure 4). In cohort 1, the most family differentiation occurred in seed length (F_59_ = 69.34, p < 0.001), area (F_59_ = 83.16, p < 0.001), and perimeter (F_59_ = 75.14, p < 0.001). In cohort 2, variation was more evenly distributed across all five traits. All traits showed significant maternal family effects in both cohorts.

**Figure 1:**
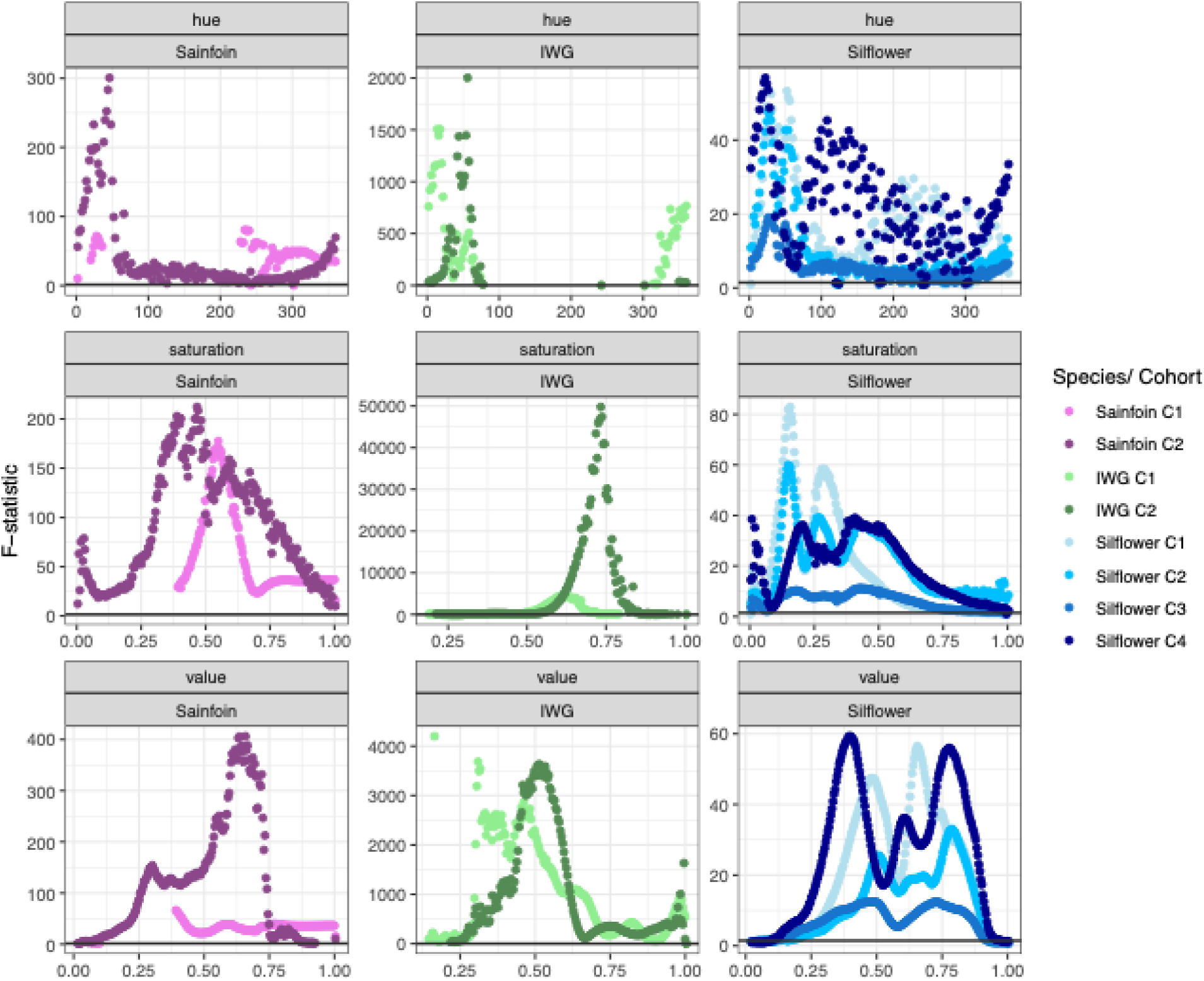
Maternal family differentiation in HSV features. Each HSV feature (hue, saturation, value) was modeled as a function of maternal family (feature ∼ family), separately by species and cohort. Features were scaled to mean = 0 and SD = 1 prior to analysis. The y-axis shows F-statistics from linear models, and the horizontal black line indicates the highest F-value that was not statistically significant (i.e., all values above the line are significant). Features with zero or near-zero variance were removed prior to modeling (see Methods). Effect sizes for each feature are shown in Supplemental Figure 1.

**Figure 2.**
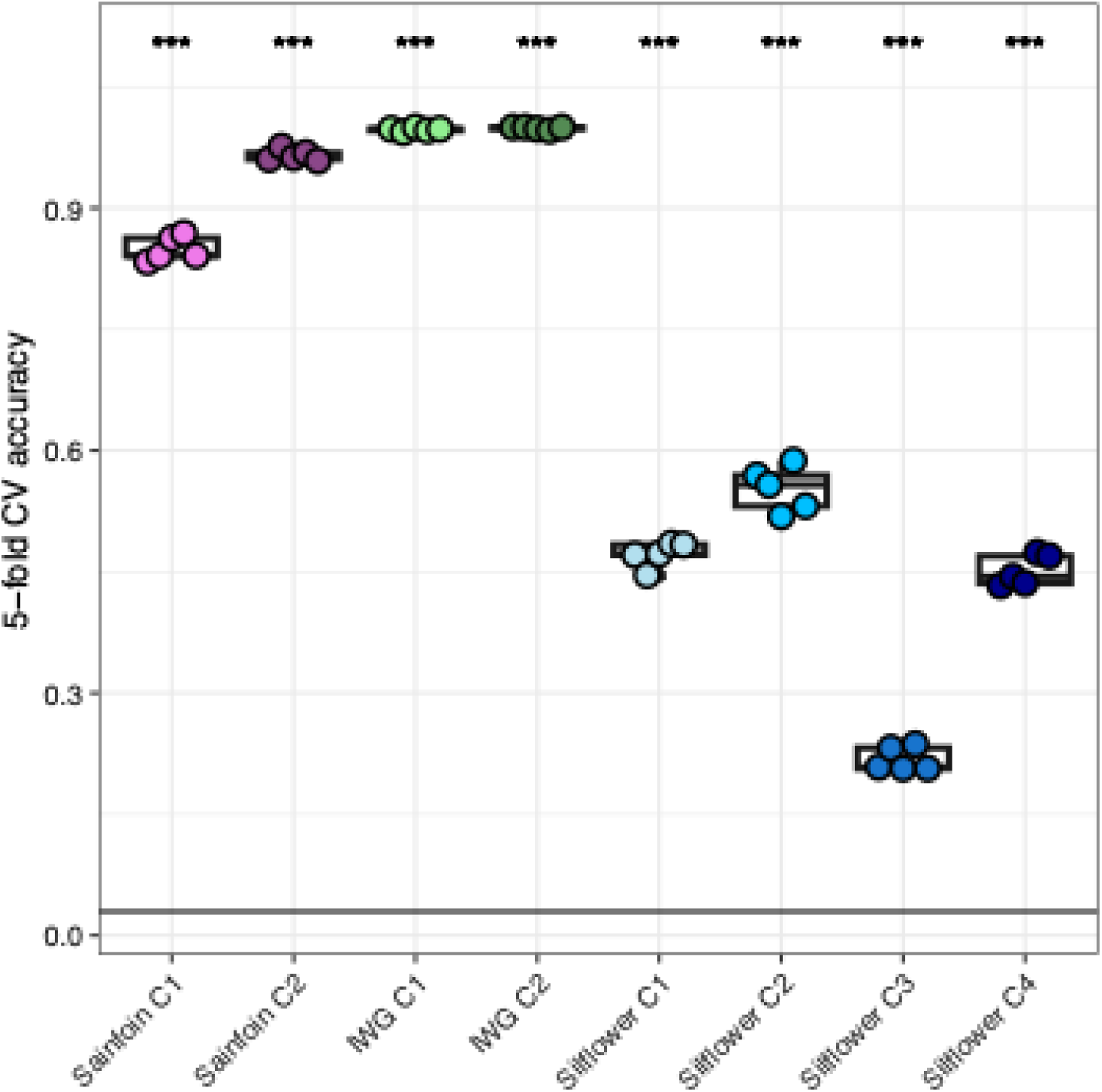
Seed HSV features predict maternal family. Across species and cohorts, seed HSV features accurately predict maternal family identity. Each model was fit using 5-fold cross-validation: 80% of data was used for training and 20% was withheld for testing in each fold. Accuracy is reported on the y-axis. Asterisks (***) indicate performance significantly better than a naïve classifier that always predicts the most frequent family (p < 0.001). Feature importance values from these models are shown in Supplemental Figure 2.

**Figure 3.**
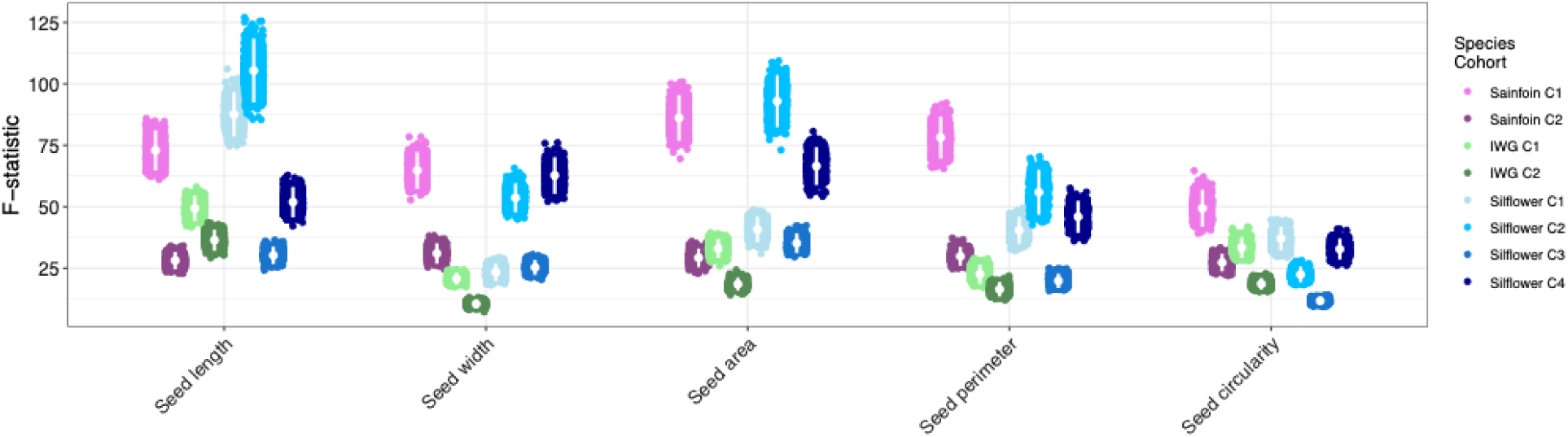
Maternal family differentiation in seed morphology. Seed morphological traits vary significantly by maternal family across species and cohorts. For each trait, we calculated the F-statistic from linear models (trait ∼ family) using 1000 bootstrap resamples. Points represent mean F-statistics; error bars show ±1.96 standard deviations. Bootstrapped estimates of variance explained (PVE) for each trait are shown in Supplemental Figure 4.

Maternal families showed significant stratification across nearly all HSV features in IWG (Figure 1). Maternal family explained the most hue variation at features <100° in both cohorts, with the strongest effects observed at 18° in cohort 1 and 56° in cohort 2. Across both cohorts, more than 75% of variation in many saturation and value features was attributable to maternal family, with strong peaks in cohort 2 at saturation ∼0.75 and value ∼0.55. LightGBM prediction accuracy of maternal family was 99.7% and 99.9% in cohorts 1 and 2, respectively (both p < 0.001; Figure 2). Morphological variation was also variable across maternal families (Figure 3; Supplemental Figure 4). In both cohorts, seed length showed the highest maternal family differentiation (F_94_ = 46.35 and F_95_ = 34.09, respectively, both p < 0.01). All morphology traits were significantly structured by maternal family.

Because our IWG samples came from a partially sequenced breeding pool, we also tested whether HSV variation was attributable to paternal family. Paternal family explained far less phenotypic variation than maternal family, and the most informative features varied by cohort (Supplemental Figure 5). In hue, variation due to paternal family in cohort 1 peaked at ∼60°, while cohort 2 peaked at ∼18°. Peak saturation differentiation occurred at saturation = 0.80 in cohort 1 and ∼0.20 in cohort 2. Value features showed weak and inconsistent patterns. LightGBM assigned samples to their paternal family with accuracies of 5.6% and 8.1% in cohorts 1 and 2, respectively. Although the accuracies of these models are low, both models performed significantly better than null expectations (p < 0.05 and p < 0.01, respectively). Paternal family only weakly described patterns of morphological variation (Supplemental Figure 6)

In silflower, maternal family explained significant HSV variation across all four cohorts (Figure 1). For hue, all cohorts showed peaks below 100°, with additional peaks at 228° in cohort 1 and 108° in cohort 4. Saturation and value features also showed broad maternal family differentiation, notably at saturation ∼0.16 (cohort 1) and value ∼0.40 and ∼0.78 (cohort 4). However, maternal family prediction accuracy using HSV features was lower in silflower than in the other species, ranging from 22% (cohort 3) to 55% (cohort 2) (Figure 2). Cohort 2 consistently exhibited the clearest family structure. Among morphological traits, seed circularity showed the strongest family differentiation in cohorts 1 and 4 (Figure 3). All traits varied significantly by maternal family across cohorts.

### Variation in germination traits

Days to germination and germination proportion were significantly variable across maternal families in all cohorts for all three species (Supplemental Table 1). The mean days to germination across maternal families were 3.2 to 5.3 days in sainfoin (cohort 1 F_59_ = 3.52, p < 0.001; cohort 2 F_63_ = 3.44, p < 0.001), 2.2 to 4.0 days in IWG (cohort 1 F_94_ = 2.60, p < 0.001; cohort 2 F_95_ = 2.32, p < 0.001), and 5.0 to 14.4 days for silflower (cohort 1 F_30_ = 8.68, p < 0.001; cohort 2 F_61_ = 11.44, p < 0.001; cohort 3 F_141_ = 4.10, p < 0.001; cohort 4 F_43_ = 15.90, p < 0.001). Sainfoin germination proportion ranged from 23.8% to 100% across maternal families (cohort 1 χ^2^_59_ = 342.53, p < 0.01; cohort 2 χ^2^_63_ = 387.16, p < 0.001). The maternal family germination proportion ranged from 64.0% to 100% for IWG (cohort 1 χ^2^ _94_ = 159.2, p < 0.001; cohort 2 χ^2^ _95_ = 155.41, p < 0.001). Silflower had maternal family germination proportions ranging from 0.8% to 100% (cohort 1 χ^2^ _30_ = 260.90, p < 0.001; cohort 2 χ^2^ _61_ = 819.29, p < 0.001; cohort 3 χ^2^ _141_ = 765.35, p < 0.001; cohort 4 χ^2^_43_= 524.22, p < 0.001).

### Correlation between seed morphological traits, HSV, and germination

Before building predictive models, we first tested whether any seed morphological traits reliably correlated with either days to germination or germination proportion across maternal families. In sainfoin cohort 1, we found that days to germination correlated with seed length (r = 0.42, p < 0.001), width (r = 0.40, p < 0.001), area (r = 0.43, p < 0.001), and perimeter (r = 0.41, p < 0.001), but these were not found in cohort 2 (Supplemental Figure 7. In IWG cohort 2, we found correlations between day to germination and seed width (r = 0.27, p < 0.01), seed area (r = 0.32, p < 0.01), and seed perimeter (r = 0.30, p < 0.01), but none of these correlations were observed in cohort 1 (Supplemental Figure 8). In IWG cohort 1, we found significant correlations between germination proportion and seed length (r = -0.30, p < 0.01), seed area (r = -0.31, p < 0.01), and seed perimeter (r = -0.27, p < 0.01), none of which were observed in cohort 2. Finally, in silflower, we observed significant correlations between days to germination and seed length (r = -0.46, p < 0.01), seed area (r = -0.49, p < 0.01), and seed perimeter (r = -0.49, p < 0.01), none of which were observed in cohorts 2-4 (Supplemental Figure 9).

To explore the utility of HSV features in predictive models, we also evaluated correlations of HSV with germination traits. We found that HSV features correlate with germination, though these relationships are often modest and vary across species and cohorts (Supplemental Figure 10). For example, in sainfoin cohort 2, we observed consistent negative correlations (r < -0.3) between germination proportion and hue features > 60° that are not observed in cohort 1. However, both cohorts demonstrate negative correlations (r < -0.25) between germination proportion and saturation and value features > 0.75. Days to germination has fewer correlates with HSV features across species, but significant correlations were identified in IWG and silflower. These patterns of correlation support using methodologies like phenomic selection to move away from feature-by-feature attempts to identify models with predictive value toward models that consider the total landscape of HSV features at one time.

### Predicting germination traits from phenomic selection models

We found that PS models incorporating relationship matrices derived from seed color features (H, S, and/or V) and morphological traits (Morph) can predict germination proportion and timing (Figure 4; Supplemental Table 2; Supplemental Table 3). The prediction accuracies vary based on the relationship matrix used in the PS model, the trait being predicted, the species, and the cohort within species (Table 3, S15). For both germination traits, each species cohort has at least one matrix predicting significantly above *r* = 0. HSVMorph was typically the best-performing matrix for days to germination across both species and cohorts, notably outperforming morphology alone.

**Figure 4.**
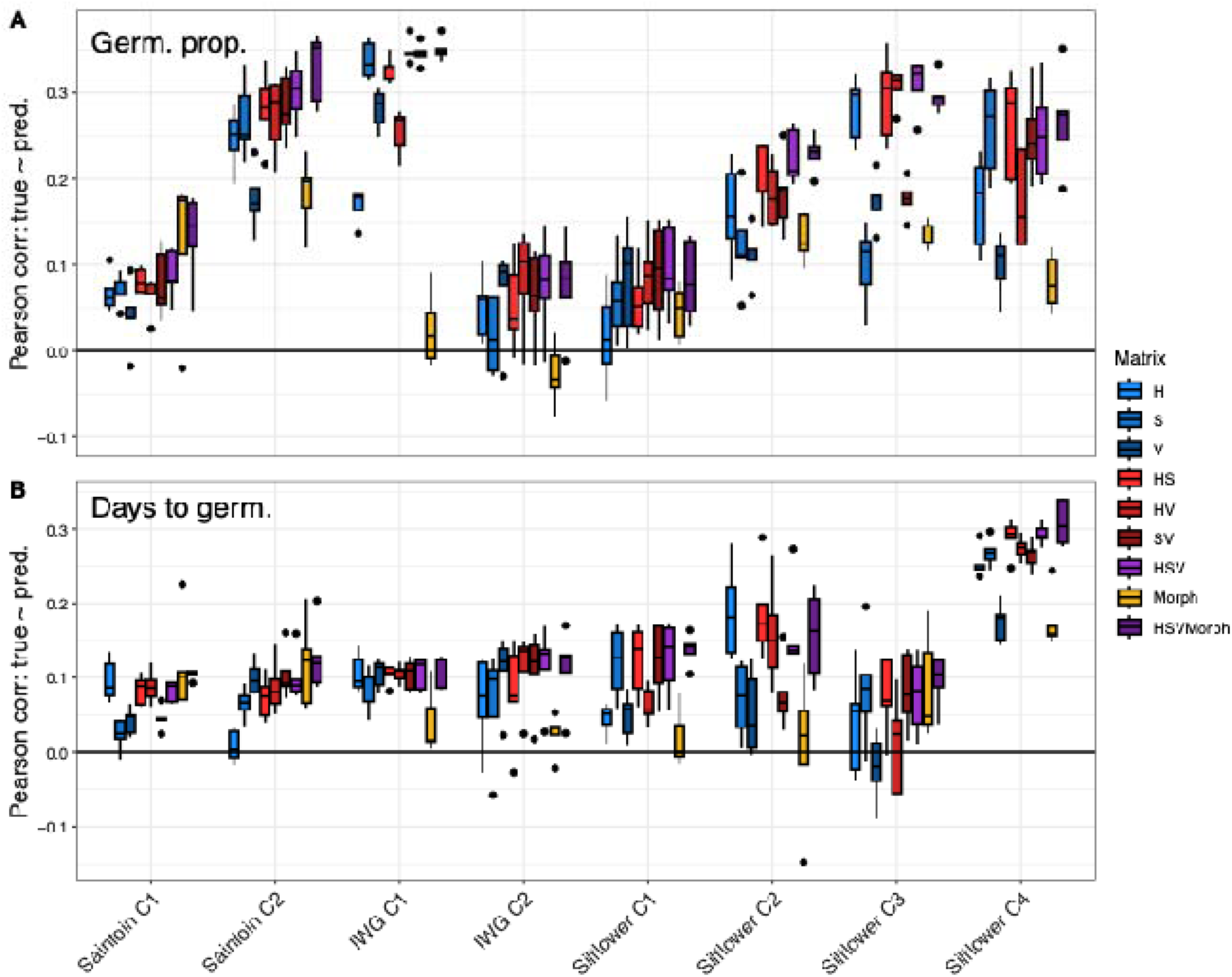
Prediction of germination traits using phenomic selection models. Phenomic selection models predict (A) germination proportion and (B) days to germination with varying accuracy depending on species, cohort, and input features. Each model was parameterized using a relationship matrix derived from one of nine data sets: hue (H), saturation (S), value (V), all combinations of HSV, seed morphology (Morph), and HSV + morphology (HSVMorph). Models based on single feature sets are shown in blue, pairwise combinations in red, and saturated models (HSV or HSVMorph) in purple. Each model was evaluated using 5-fold cross-validation, and the y-axis shows Pearson’s correlation (r) between predicted and observed trait values in the test set.

In sainfoin cohort 1, the highest average performing matrices to predict germination proportion were Morph and HSVMorph (r = 0.13), though they did not statistically differ from any other matrix except V. For days to germination, Morph alone yielded the highest average correlation (r = 0.11), but was equivalent to all matrices except S and V. In cohort 2, predictive performance improved. HSVMorph provided the strongest prediction for germination proportion (r = 0.33) and was statistically distinct from Morph alone (r = 0.18). For days to germination in cohort 2, HSVMorph had the highest performance (r = 0.10), but was statistically indistinguishable from Morph and HSV alone.

In IWG cohort 1, several matrices achieved an average performance r ∼= 0.35 in predicting germination proportion. Notably, S, SV, HSV, and HSVMorph all outperformed Morph alone. Similarly, days to germination saw better performance in all matrices as compared to Morph alone, but these performances were much smaller (r = 0.1) and not statistically significant. In cohort 2, performance dropped considerably for germination proportion, with the best performance model (V) outperforming only Morph alone. Performance for days to germination was similar to cohort 1, with HSV and HSVmorph both statistically outperforming morph alone.

In silflower cohort 1, all matrices performed equivalently in predicting germination proportion, with HSV yielding the highest average correlation (r = 0.10). For days to germination, both HSV and HSVMorph were the best predictors (r = 0.14), and performed statistically better than Morph alone. In cohort 2, HSV and HSVMorph again performed best (r = 0.23 for germination proportion), with both matrices showing statistical differentiation from Morph. Cohort 3 exhibited the highest germination proportion predictions across all silflower cohorts with HSV peaking at r = 0.31, far outperforming Morph alone. However, predictions for days to germination in this cohort were weaker, with HSVMorph leading at r = 0.10, though not distinguishable from any other matrix. In cohort 4, HSV and HSVMorph were the top predictors for germination proportion (r = 0.27) and for days to germination (r = 0.31), significantly outperforming other matrices, especially V and Morph.

## Discussion

There is growing interest in the development of new crops that simultaneously meet the needs of humans and provide ecosystem services like the prevention of soil erosion, improved nutrient capture, and enhanced carbon sequestration (Griffiths et al., 2022; Jackson, 2002; Rapidel et al., 2015). Some of the systems that are most likely to provide these services, herbaceous perennial plants, are underrepresented in agriculture (Van Tassel et al., 2010). To overcome the paucity of herbaceous perennial systems in modern agriculture, rapid development of perennial counterparts to large industrialized cropping systems is necessary. We hope to leapfrog both the 10,000 years of selection and classic plant breeding, domesticating new perennial crops using technological innovations such as phenomic selection that predicts primary traits with cheaper, easy-to-measure secondary traits (Rincent et al., 2018; Van Tassel et al., 2022).

In this study, we showed that high-dimensional seed color profiles can improve the prediction of key germination traits within a PS framework – traits that must be stabilized and optimized during the domestication of novel crops. We demonstrated that predictions are possible due to extensive variation in both secondary traits (color and morphology) and primary traits (germination proportion and days to germination). We detailed the improvements of prediction when high-dimensional color traits are used to either replace or supplement seed morphological data. Finally, we examined which features or combinations of features within the high-dimensional colorspace provide the most information for predicting germination traits. This work adds to an expanding body of literature that supports the potential of high-dimensional phenotypes in breeding pipelines and sets the stage for future work predicting agronomic traits in the field using phenomic selection.

### High-dimensional secondary traits predict maternal identity

Given the extensive variation in seed colorspace and morphological traits attributable to maternal family, we assessed whether these differences arise from either genetic or non-genetic maternal effects. While we were unable to test this in sainfoin and silflower, a subset of individuals in the IWG experiment have been genotyped, and as such, we could identify the paternal parent of each seed. In all traits measured, we identified that the effect of the paternal family was considerably smaller than the maternal parent. This is not unexpected, as maternal effects have significant impacts on seed development and dormancy (Postma & Ågren, 2015; Roach & Wulff, 1987). The maternal environment contributes to offspring germination success and transgenerational phenotypic plasticity (Lacey, 1996; Schmid & Dolt, 1994). The presence of non-genetic maternal effects and their contribution to variation in key germination traits does not undermine the potential for selection; rather, it highlights the importance of accounting for these effects in both interpretation and experimental design.

Significant correlations among morphological traits were also present in all species and cohorts, but correlations with germination traits were rare and inconsistent across cohorts, which comports with other studies of germination in perennial species (Herron et al., 2020, 2021). This suggests that morphology alone is not a reliable indicator of germination in these species. This is a contentious point in ecological theory (Westoby et al., 1992), which has historically reported relationships between seed size and germination (Aiken & Springer, 1995; Bond et al., 1999; Harper et al., 1970). However, if morphology alone lacks a relationship to germination traits, other phenotypes might be more useful, either alone or in supplement to morphology in developing predictive models.

### High-dimensional relationship matrices predict germination

In this study, relationship matrices from high-dimensional HSV color data and morphology traits show promise in predicting germination proportion and days to germination. Given the extensive variation in HSV color traits and seed morphological traits, we expected that expanding the feature space, both by including more HSV dimensions and incorporating morphology, would enhance phenomic selection performance. While HSV features capture detailed variation in pigmentation, morphology offers complementary structural information about seed size and shape, which may further strengthen model predictions. Consistent with these expectations, the trait matrix with the highest dimensionality (HSV or HSVMorph) generally yielded the most accurate predictions.

Predicting whether a seed will germinate and increasing the overall germination proportion in new crops is important for both breeders and farmers alike. Germination occurs as seeds shift from a dormant state through emergence and into active growth as a seedling. Wild plant species exhibit varying germination strategies. In some species, germination requires physical or chemical stimulus (Nelson et al., 2012; Olszewski et al., 2010), some seeds must experience cold to germinate (Bratcher et al., 1993), and yet others exhibit staggering, where seedlings emerge at different times during a season (Mercer et al., 2011) or across multiple years (Cohen, 1966). In domesticated species, this behavior, called bet-hedging, has been selected against during the domestication process, as most industrialized agricultural systems require that seeds germinate in synchronicity (Finch-Savage & Bassel, 2016). Variation in seed traits affects germination success, and this variation often arises from differences across populations (McWilliams et al., 1968). Historical selection based on seed size alone has shown a correlation between seed size and yield, and that seed size is a heritable trait (Harper et al., 1970). The ability to predict days to germination is an important step in breeding toward synchronizing germination.

Seed color analysis using RGB values has been used to predict germination in crops like rice (Lurstwut & Pornpanomchai, 2017), castor (Moosavi et al., 2022), and perennial crops sainfoin and alfalfa (Behtari et al., 2014). Moreover, RGB-deep learning models have been developed to predict germination in discarded produce seed lots (Nehoshtan et al., 2021). The ability to predict germination using models derived from seed color is supported by the underlying biological processes within a seed. In sheepgrass, anthocyanin/proanthocyanidin-specific pathway genes, which control seed coat color, have been correlated to dormancy strength (Zhao et al., 2019). Legume germination is controlled by temperature changes and increased seed coat permeability, the strength of which is correlated to coat color (Smýkal et al., 2014). To our knowledge, this paper is the first to demonstrate the use of high-dimensional seed color data (HSV) as a method to predict traits using phenomic selection in perennial crops.

In addition to visible light, interaction with light in the NIR region has been successfully used as the basis for PS models. For example, Graciano et al. (2025) used NIR reflectance from individual sweet corn kernels to successfully predict germination with a Pearson’s coefficient of *r* = 0.58, and Lane et al. (2020) used NIR from maize kernels to predict yield up to *r* = 0.74. Similarly, Laurençon et al. (2024) predicted germination using NIR data from oilseed rape dry seeds at approximately *r* = 0.3, and germination time around *r* = 0.5. This study demonstrated consistent predictive ability, even when training data was taken from seeds harvested from a different location than where the individuals being predicted were grown. Future work should be done to determine if NIR reflectance holds similar predictive capacity in our species and whether supplementing with high-dimensional HSV traits and/or morphology can yield even better predictions.

Our predictions of germination traits using seed HSV and morphology traits represent a promising approach to accelerate the development of emerging crops. Prediction accuracies compare favorably to prior work using spectral reflectance in well-studied systems such as rapeseed (Laurençon et al., 2024). A key innovation of this work lies in the use of high-dimensional phenotypes derived from simple, low-cost consumer-grade document scanners, eliminating the need for specialized hyperspectral or multispectral imaging equipment. This approach significantly lowers the technological and financial barriers to implementing PS, broadening accessibility for breeding programs with limited resources. Analyzing seed color within a high-dimensional colorspace captures variation that extends the constraints of standard RGB models and enables more nuanced models. In addition, seed traits can be collected nondestructively, allowing predictions to inform selection decisions before planting. By predicting germination outcomes and other traits from seed phenotypes, breeders can effectively screen and discard less promising lines before investing resources in planting and field evaluation, reducing time and labor requirements. Together, these advantages position this method as a scalable and practical tool for integrating phenomic data into breeding pipelines, particularly for under-resourced and emerging perennial crops.

### Limitations

While our results demonstrate the potential for using image-derived seed traits in phenomic selection, several limitations should be acknowledged. First, although some models achieved modest predictive accuracy (e.g., *r* = 0.33), others performed closer to baseline levels. These modest correlations, especially for days to germination, reflect the complexity of these traits and the likelihood of noise, both biological and technical.

A second limitation is the absence of genomic data for most individuals in this study. While we observed strong maternal family effects, we were unable to fully disentangle genetic from non-genetic maternal influences, which are known to affect early life traits like germination and dormancy. Moreover, we were unable to compare phenomic predictions from seed color to GS methods. Although a subset of individuals in this study were genotyped, our focus was specifically on evaluating whether seed-based phenotypes could be used for early-stage prediction. In the case of IWG, DNA extraction was from plant tissue and not seeds, as doing so would have precluded evaluating the germination traits central to this work.

Third, although our use of HSV color features and morphology presents a cost-effective, high-throughput alternative to hyperspectral imaging, these image-derived features may not capture all relevant biochemical variation. Incorporating additional data types, such as seedling or leaf NIR reflectance, would increase the dimensionality used to compute the relationship matrices and could improve prediction accuracy even further. Post-germination traits, including biomass, overwinter survival, and yield, may also be predictable from seed traits, but were not assessed in this study.

Finally, generalizing PS models across spatial and temporal gradients remains a key challenge. Our models were developed and validated within cohorts. We did not test the extent to which prediction models generalize across cohorts, environments, or years—a critical step for PS to be robust in real-world breeding pipelines. Moreover, our study contained individuals which were exposed to different seedling treatments and different transplant sites. Future analyses will test whether models trained on one set of families or environments can accurately predict outcomes in others. Such transferability is essential for PS to accelerate perennial crop breeding at scale.

## Conclusions

Developing new crops that are both productive and ecologically sustainable is a major challenge, especially for herbaceous perennials that have only recently entered the domestication pipeline. In this study, we tested whether simple, low-cost seed traits derived from high-dimensional color profiles and morphological measurements could be used to predict germination outcomes in three emerging perennial crop species. We found that these traits vary strongly by maternal family and are predictive of both germination proportion and timing. Predictions were generally more accurate when all available features were included in the model combining high-dimensional seed color traits and seed morphology. Although predictive accuracy varied across species and cohorts, several models performed significantly better than chance. Our results show that image-based seed phenotypes can support early-stage selection in breeding programs, offering a scalable, cost-effective tool to accelerate the development of new perennial crops. This approach reduces the need for expensive equipment or multi-year field trials and can help breeders focus resources on the most promising seeds before planting.

## Author Contributions (CRediT)

LRD, BS, DLVT, MKT, MJR, and AJM contributed to the conceptualization and funding acquisition of this study. WK, ZNH, JB, EC, and MJR contributed to data curation. WK, ZNH, and MJR contributed to formal analysis. WK, ZNH, JB, EC, JG, EP, HS, JS, NF, and MJR contributed to investigations, methodology, and/or software development. The project was administered by AJM and both MJR and AJM contributed to supervision. DLVT, LRD, BS, and EC contributed resources for this project. WK and ZNH created data visualization and contributed to the first draft of this manuscript. All authors reviewed and edited the final manuscript.

## Acknowledgements

We thank the Foundation for Food and Agriculture (CA20-SS-0000000123), The Perennial Agriculture Project in conjunction with the Malone Family Land Preservation Foundation and The Land Institute, Donald Danforth Plant Science Center, The Land Institute, and Saint Louis University for funding this work. We also thank members of the Miller Lab, specifically Samantha Mazumder, Sterling Herron, and Tyler Thrash, for assistance with planting and data collection, and Danforth Center Plant Growth Facilities (RRID:SCR_024902), Danforth Center Phenotyping Facility (RRID:SCR_019049) and the Danforth Center Data Science Facility.

## Data availability

Extracted seed HSV profiles, files containing germination data and time to germination, and versions of record for all code used in this paper including Jupyter Notebooks and R scripts can be found on FigShare at: (Harris, 2025). Full resolution raw seed images are available upon request.

## Supplemental Figures

**Supplemental Figure 1.**
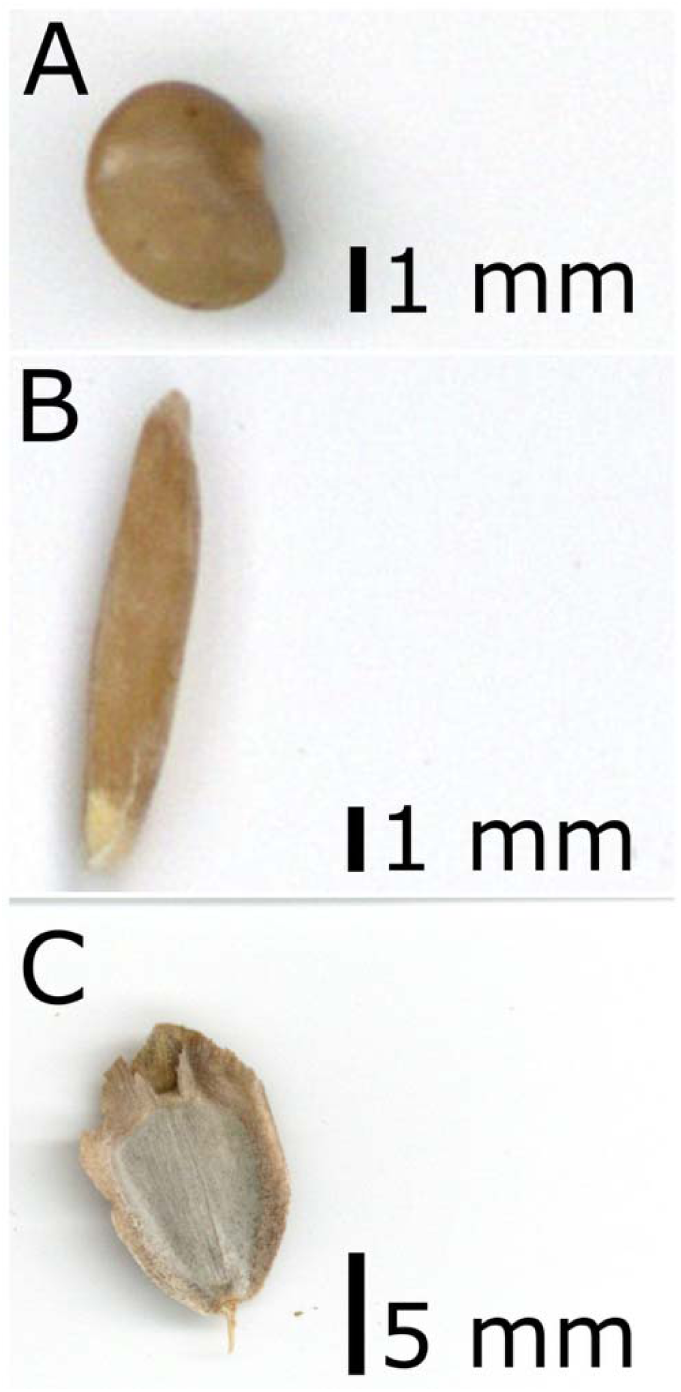
Example seed scans for each of the three species of this study: (a) sainfoin, (b) IWG, and (c) silflower.

**Supplemental Figure 2.**
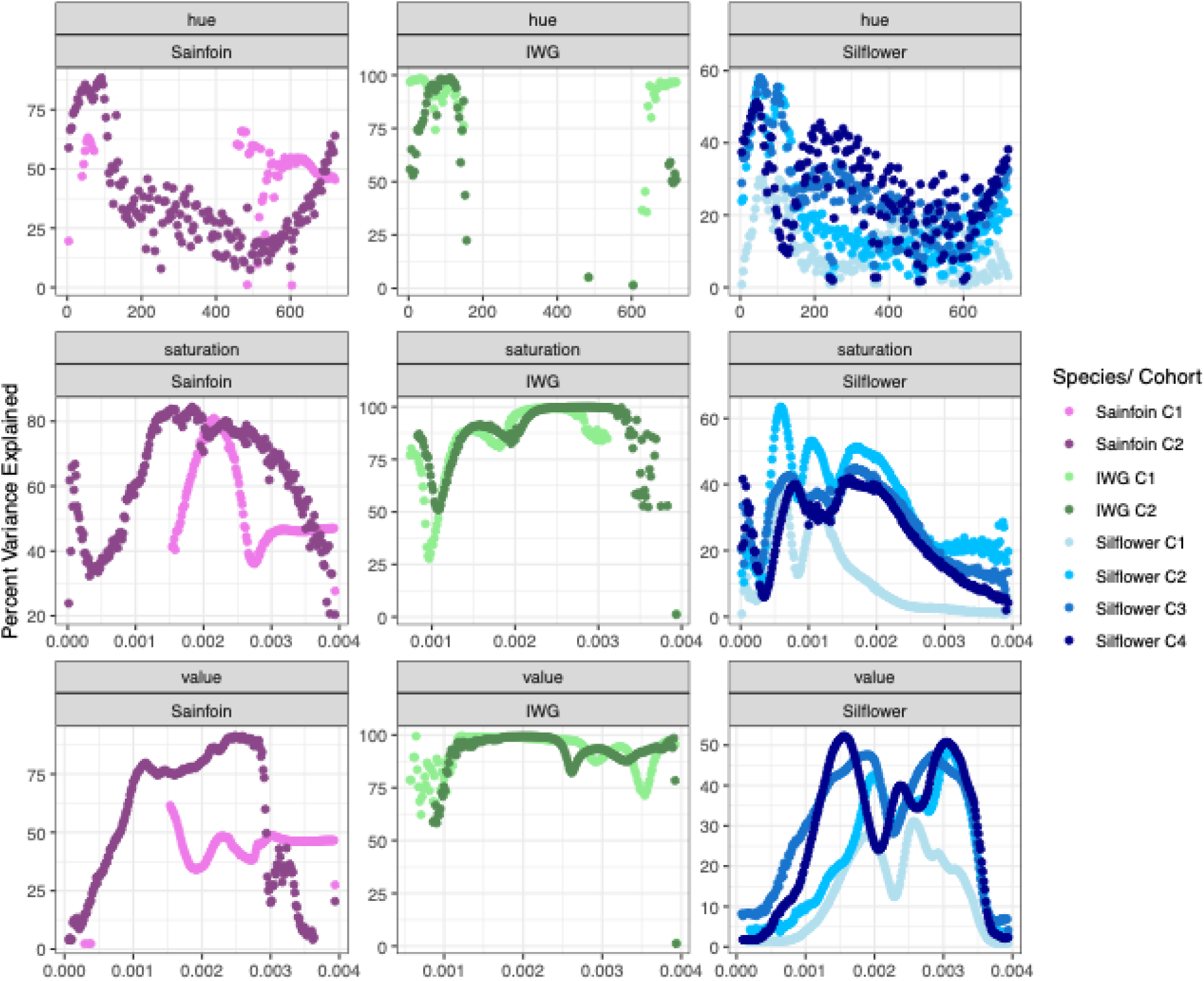
Proportion of variance explained (PVE) by maternal family in HSV features.

**Supplemental Figure 3:**
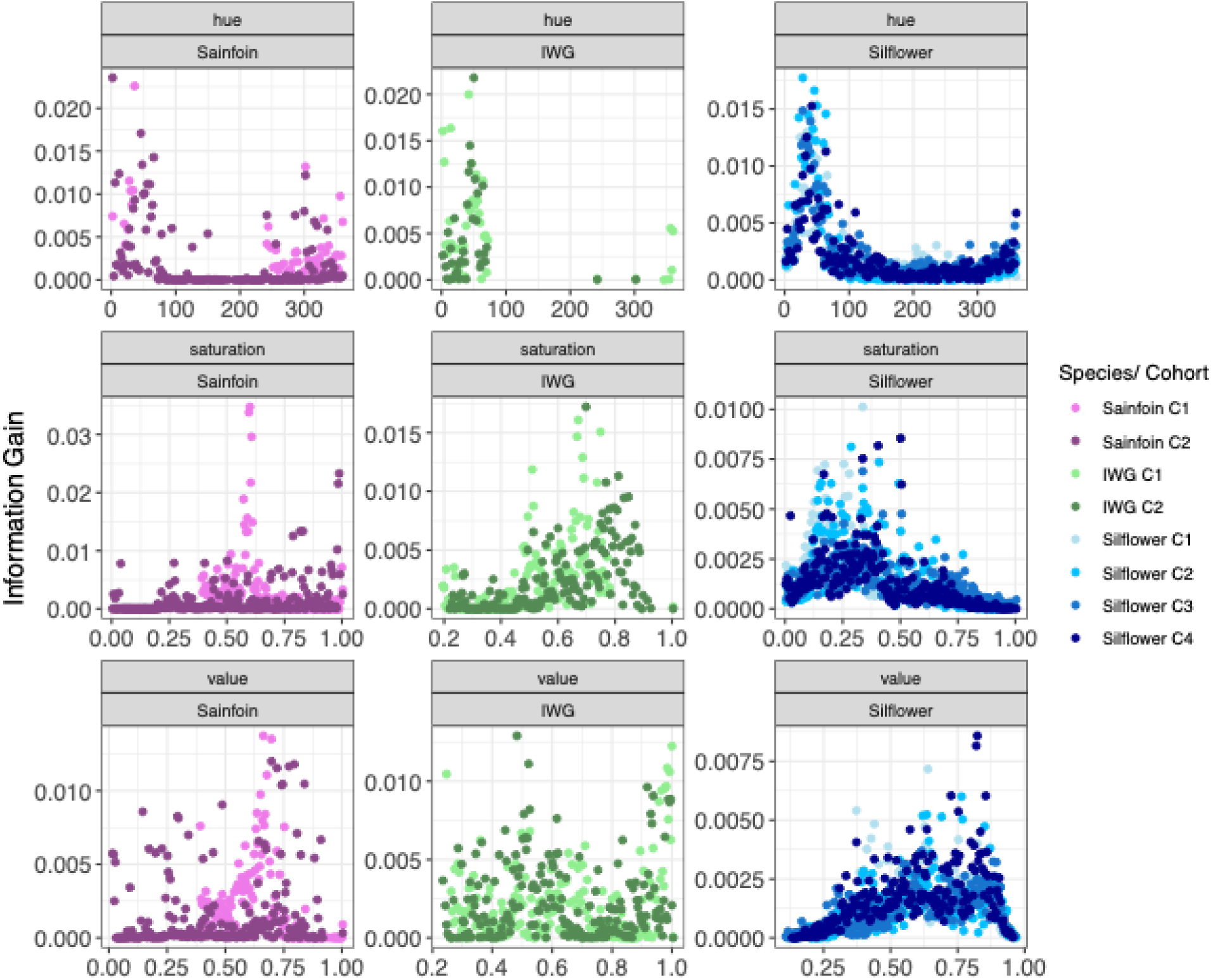
Feature importance in predicting maternal family from seed HSV

**Supplemental Figure 4:**
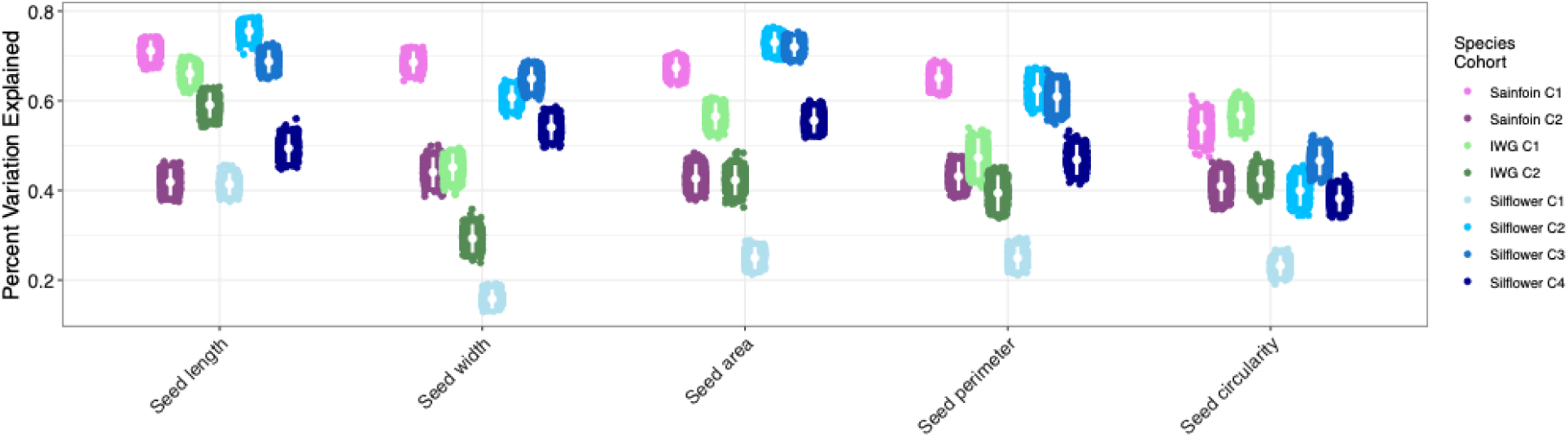
Proportion of variance explained (PVE) by maternal family in seed morphology traits.

**Supplemental Figure 5:**
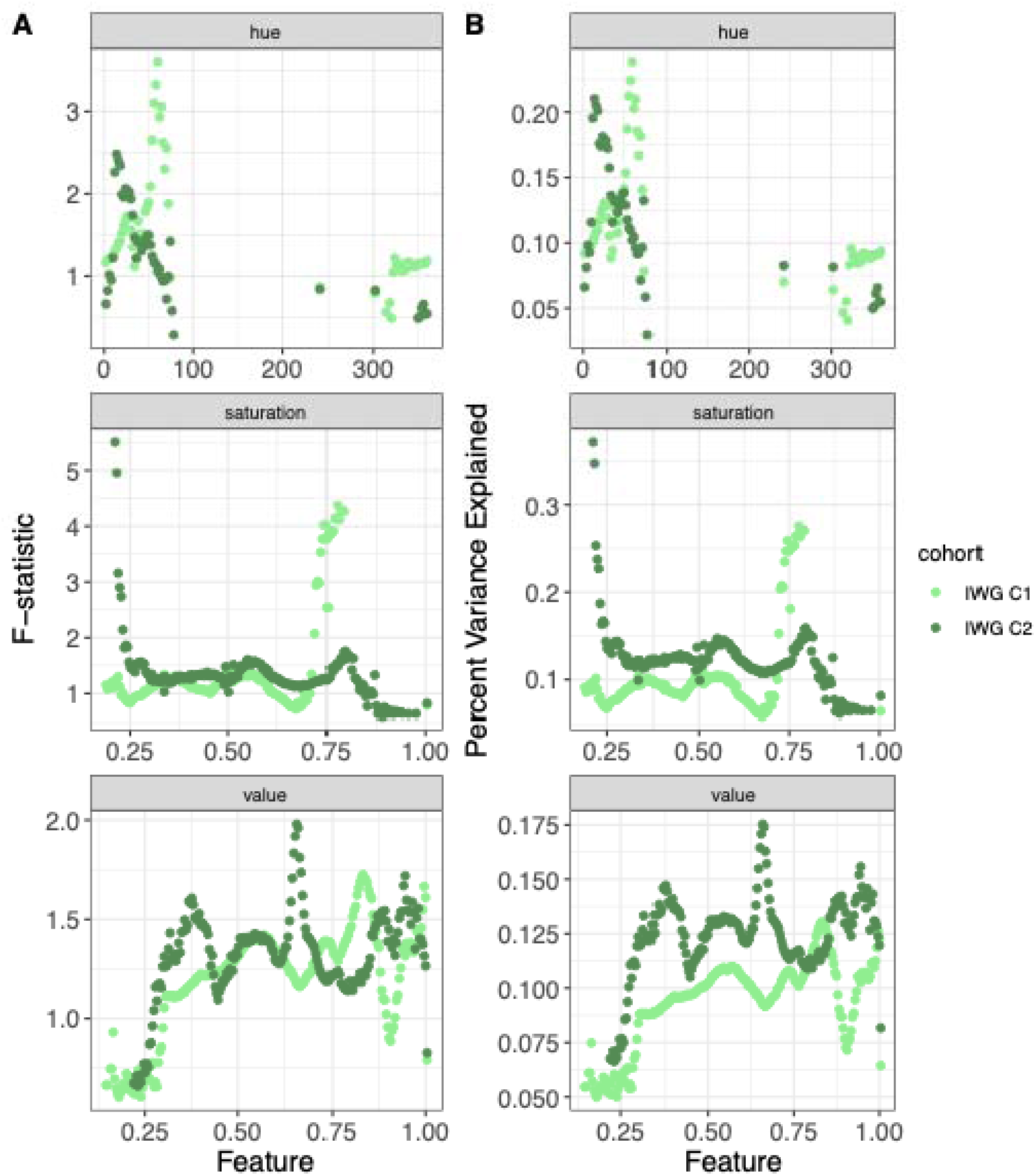
Paternal family differentiation and PVE for HSV features in intermediate wheatgrass.

**Supplemental Figure 6:**
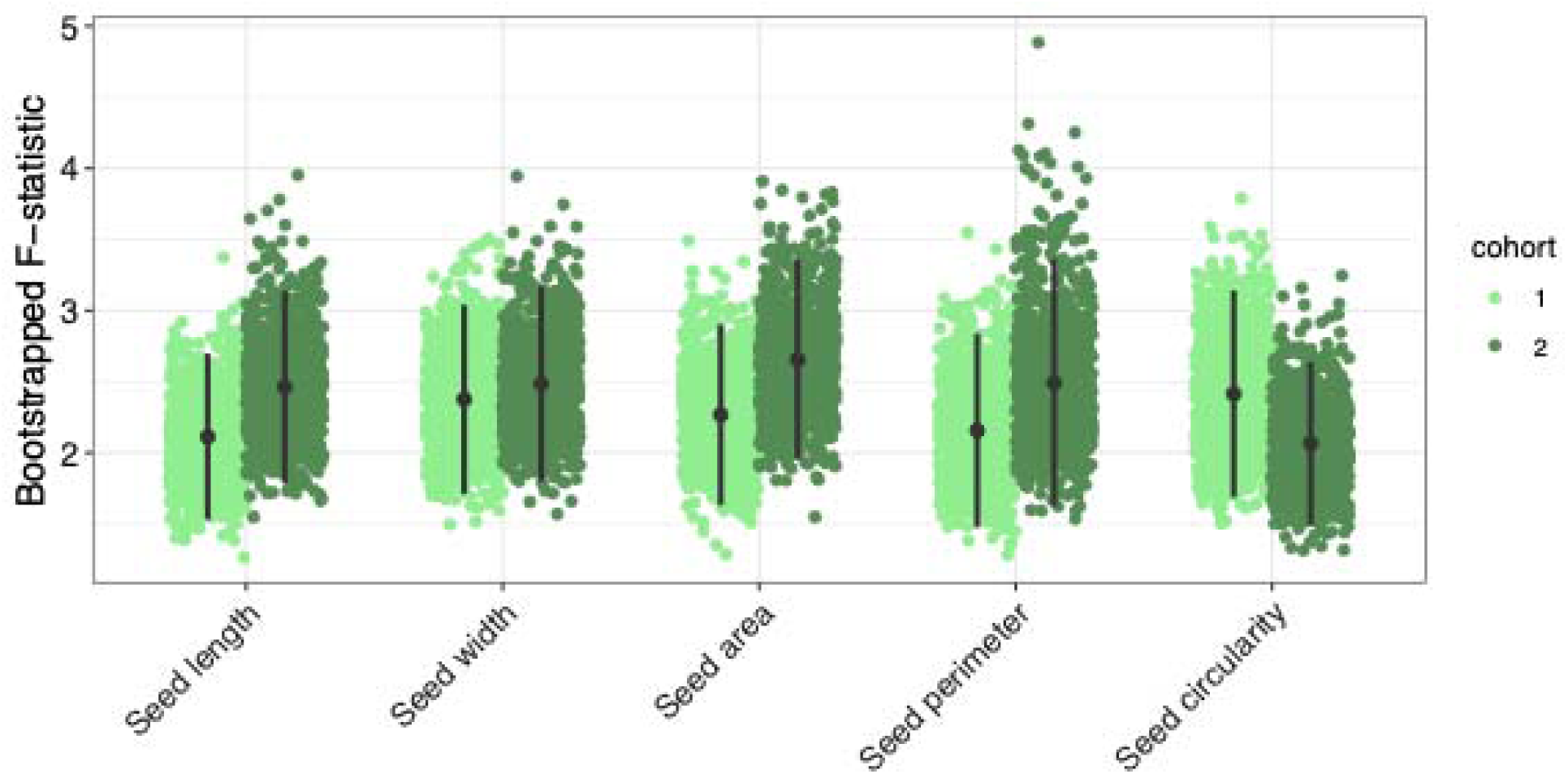
Bootstrapped F-statistics for seed morphology traits explained by paternal family in IWG.

**Supplemental Figure 7:**
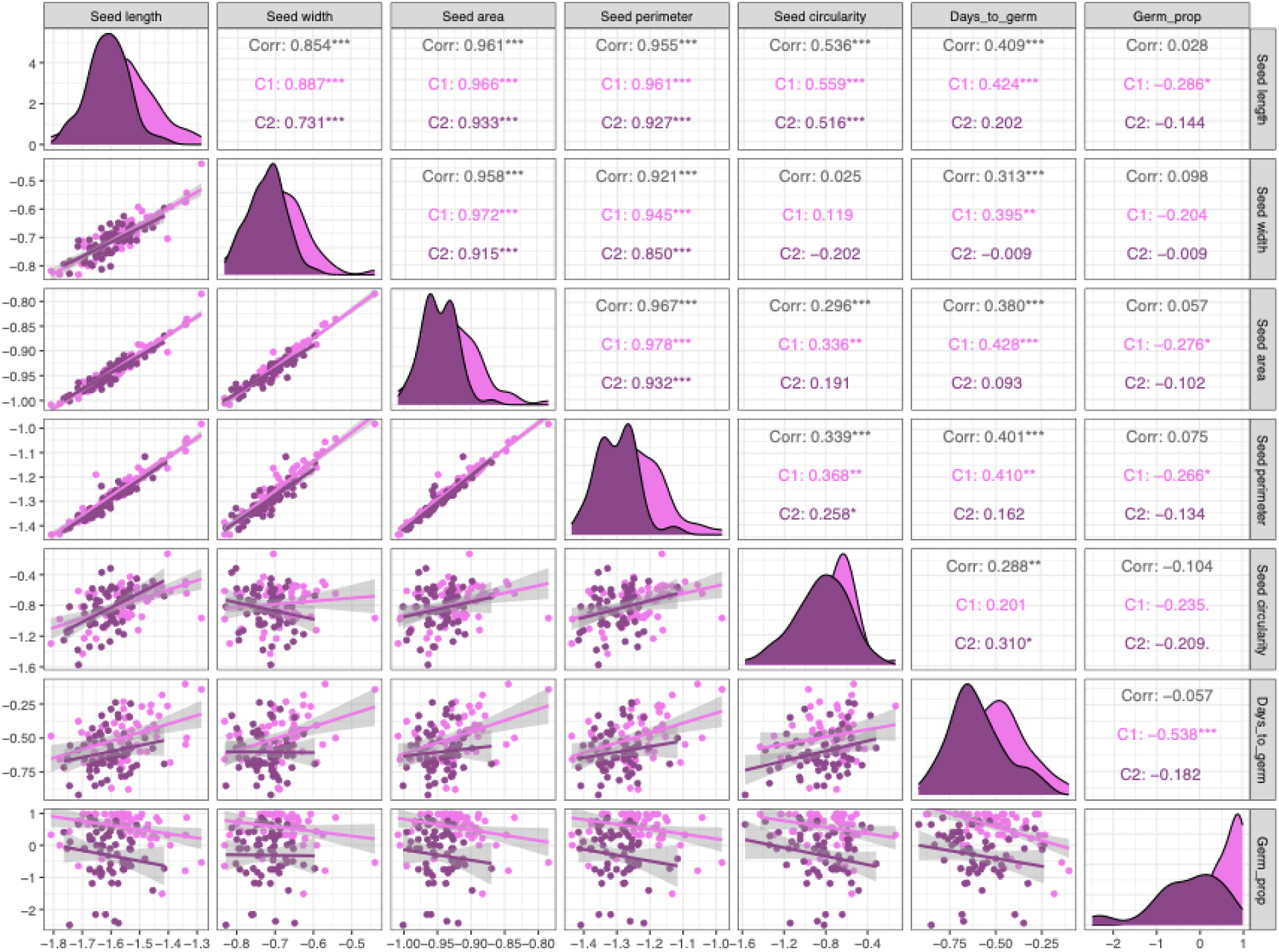
Phenotypic correlations in sainfoin. Each trait was scaled to a mean equal to 1 and unit variance across all three species. Correlations are computed by and across cohorts.

**Supplemental Figure 8:**
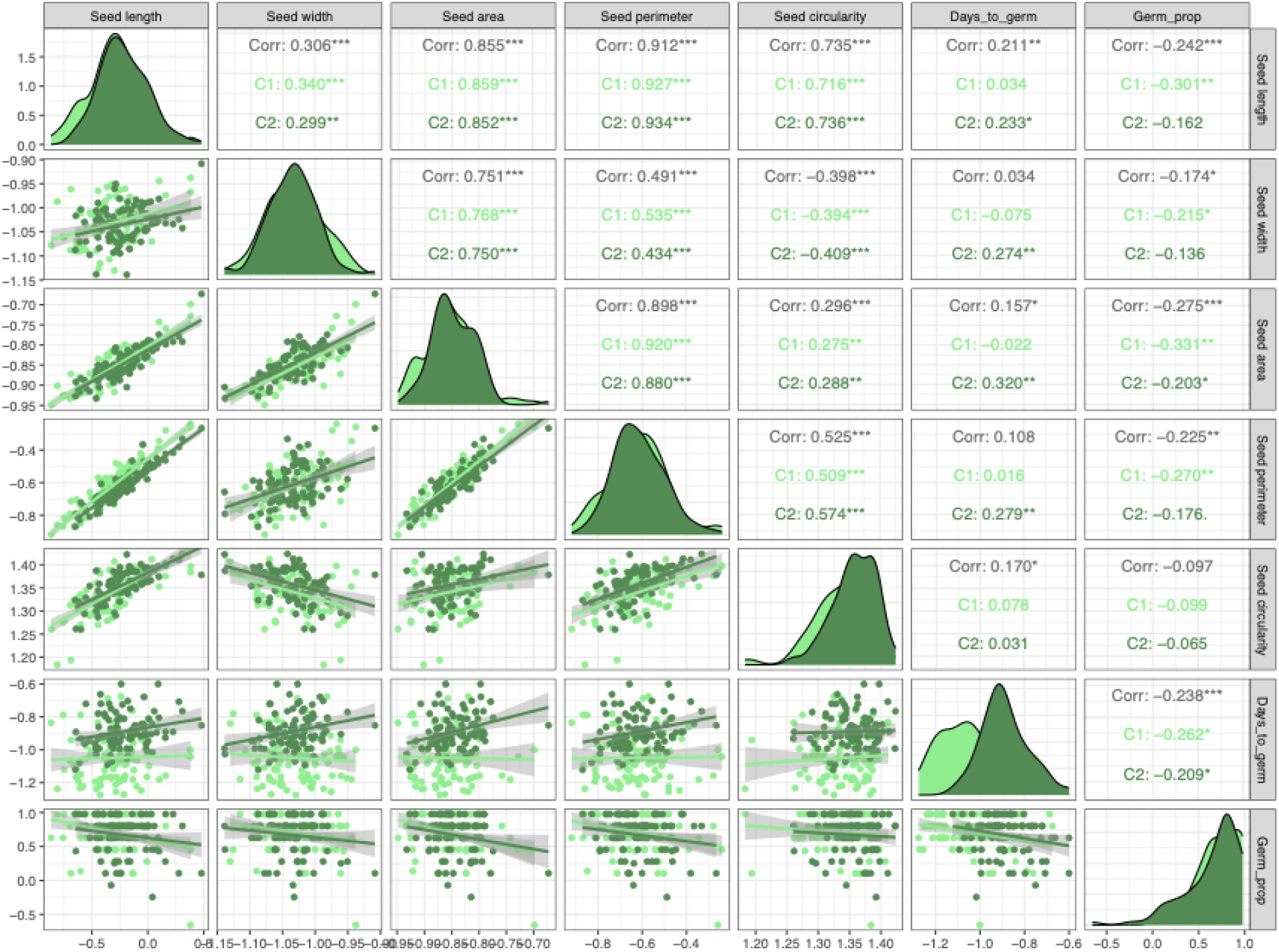
Phenotypic correlations in IWG. Each trait was scaled to a mean equal to 1 and unit variance across all three species. Correlations are computed by and across cohorts.

**Supplemental Figure 9:**
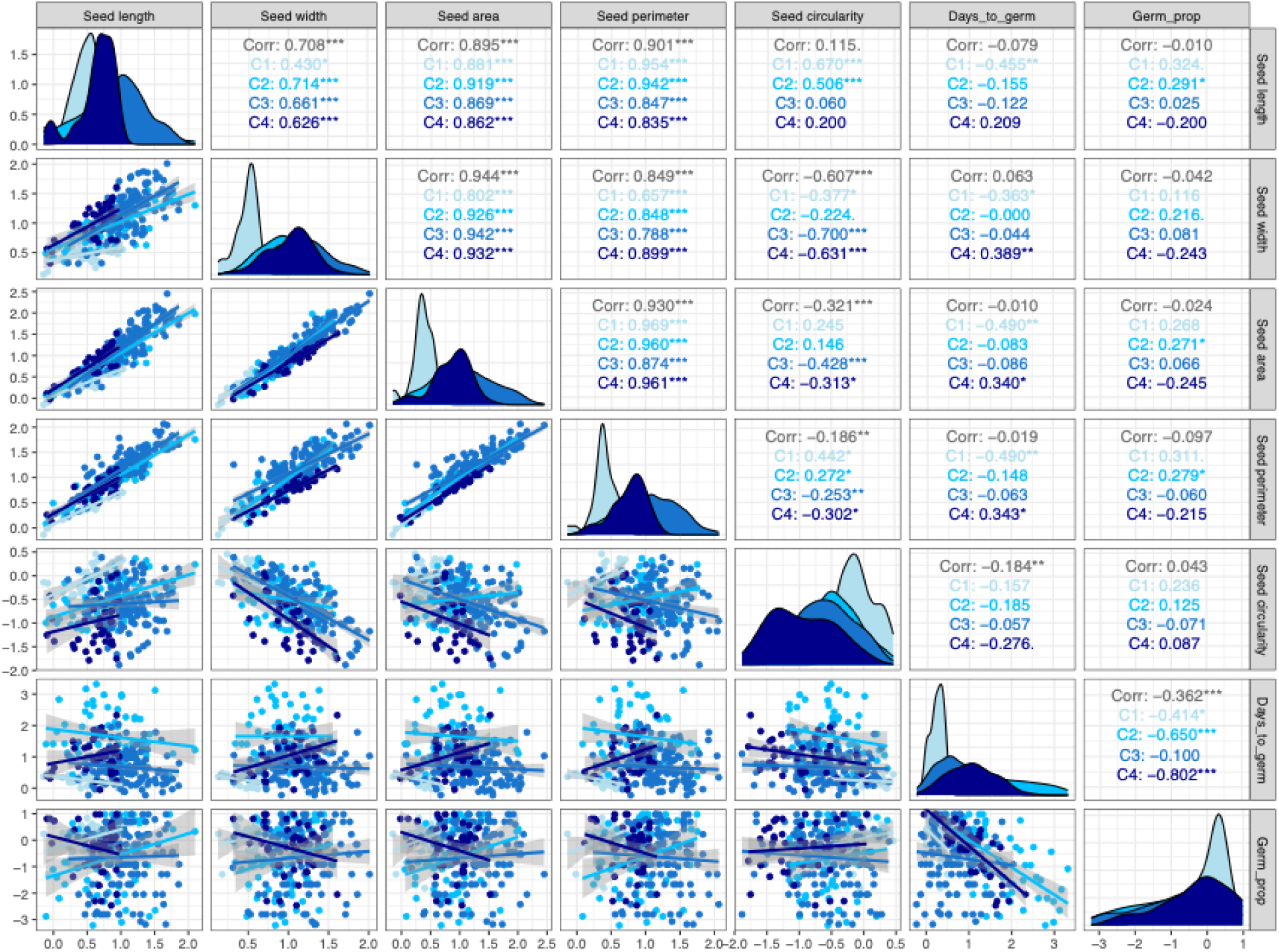
Phenotypic correlations in silflower. Each trait was scaled to a mean equal to 1 and unit variance across all three species. Correlations are computed by and across cohorts.

**Supplemental Figure 10:**
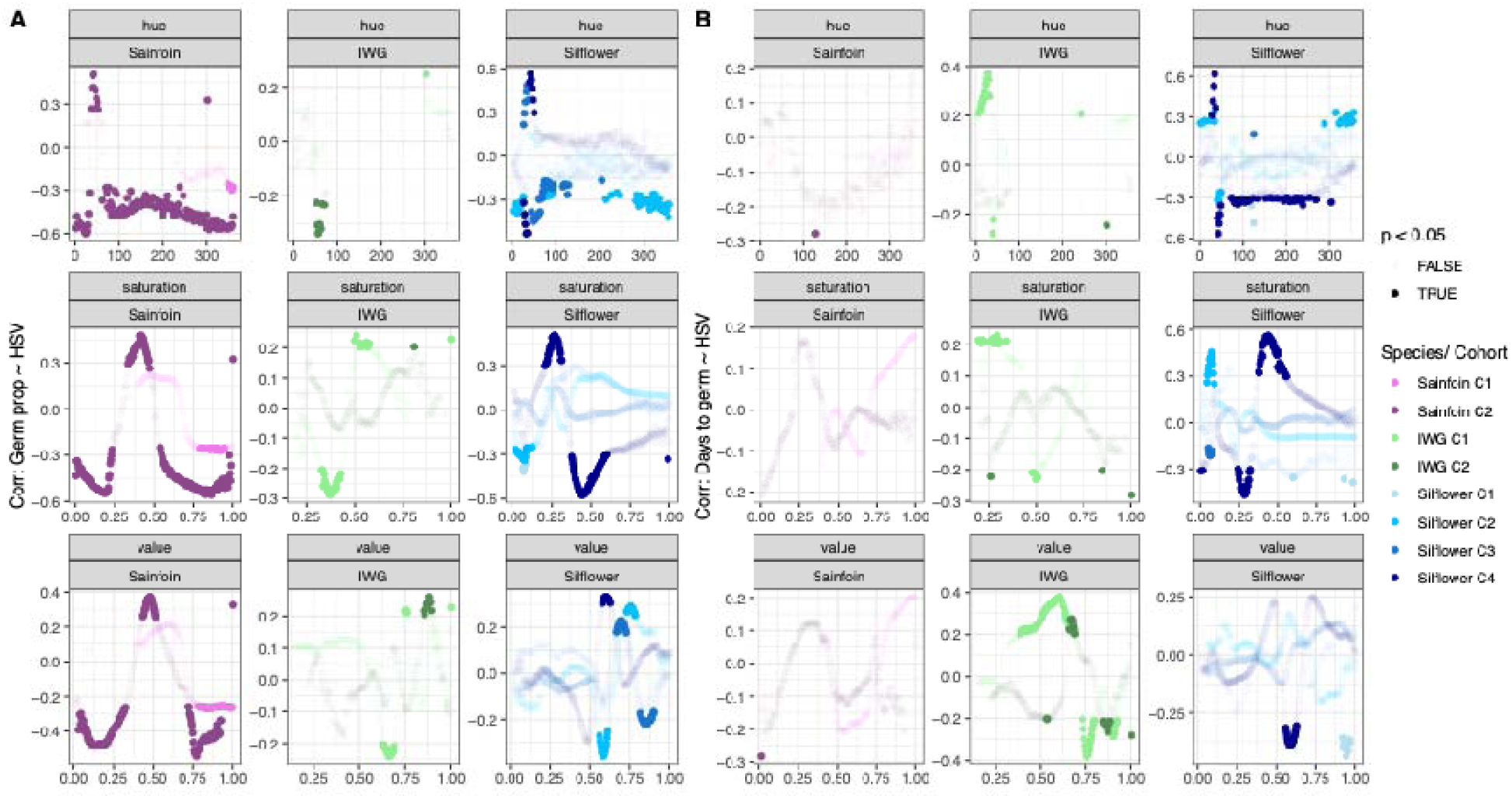
Germination trait correlations with seed HSV.

**Supplemental Table 1:** Maternal differentiation in morphological and germination traits ANOVA results for five seed morphology traits, days to germination, and germination proportion for species *Thinopyrum intermedium, Onobrychis viciifolia,* and *Silphium integrifolium*. F-statistics are reported for all traits except germination proportion where Chi-squared are reported.* P < 0.05,** P < 0.01,*** P < 0.001.

| Trait | Sainfoin C1 | Sainfoin C2 | IWG C1 | IWG C2 | Silflower C1 | Silflower C2 | Silflower C3 | Silflower C4 |
| --- | --- | --- | --- | --- | --- | --- | --- | --- |
| Seed area | $F_{59} = 83.16^{***}$ | $F_{63} = 27.52^{***}$ | $F_{94} = 30.70^{***}$ | $F_{95} = 16.85^{***}$ | $F_{30} = 39.59^{**}$ | $F_{61} = 88.55^{***}$ | $F_{146} = 31.81^{***}$ | $F_{43} = 64.10^{***}$ |
| Seed length | $F_{59} = 69.34^{***}$ | $F_{63} = 26.52^{***}$ | $F_{94} = 46.35^{***}$ | $F_{95} = 34.09^{***}$ | $F_{30} = 85.86^{***}$ | $F_{61} = 101.30^{***}$ | $F_{146} = 27.26^{***}$ | $F_{43} = 49.85^{***}$ |
| Seed width | $F_{59} = 61.59^{***}$ | $F_{63} = 29.12^{***}$ | $F_{94} = 19.04^{***}$ | $F_{95} = 9.08^{***}$ | $F_{30} = 22.22^{***}$ | $F_{61} = 50.68^{***}$ | $F_{146} = 22.65^{***}$ | $F_{43} = 60.64^{***}$ |
| Seed perimeter | $F_{59} = 75.14^{***}$ | $F_{63} = 28.07^{***}$ | $F_{94} = 20.53^{***}$ | $F_{95} = 14.81^{***}$ | $F_{30} = 39.15^{***}$ | $F_{61} = 53.30^{***}$ | $F_{146} = 17.33^{***}$ | $F_{43} = 43.91^{***}$ |
| Seed eccentricity | $F_{59} = 47.05^{***}$ | $F_{63} = 25.54^{***}$ | $F_{94} = 31.05^{***}$ | $F_{95} = 16.96^{***}$ | $F_{30} = 35.76^{***}$ | $F_{61} = 20.79^{***}$ | $F_{146} = 10.03^{***}$ | $F_{43} = 31.09^{***}$ |
| Mean hue | $F_{59} = 16.30^{***}$ | $F_{63} = 23.54^{***}$ | $F_{94} = 1.88^{***}$ | $F_{95} = 2.04^{***}$ | $F_{30} = 33.68^{***}$ | $F_{61} = 41.42^{***}$ | $F_{146} = 16.36^{***}$ | $F_{43} = 31.28^{***}$ |
| Days to germination | $F_{59} = 3.52^{***}$ | $F_{63} = 3.44^{***}$ | $F_{94} = 2.60^{***}$ | $F_{95} = 2.32^{***}$ | $F_{30} = 8.68^{***}$ | $F_{61} = 11.44^{***}$ | $F_{146} = 4.10^{***}$ | $F_{43} = 15.90^{***}$ |
| Germination Proportion | $\chi^2_{59} = 342.53^{**}$ | $\chi^2_{63} = 387.16^{***}$ | $\chi^2_{94} = 159.2^{***}$ | $\chi^2_{95} = 155.41^{***}$ | $\chi^2_{30} = 260.90^{***}$ | $\chi^2_{61} = 819.29^{***}$ | $\chi^2_{141} = 765.35^{***}$ | $\chi^2_{43} = 524.22^{***}$ |

**Supplemental Table 2:**
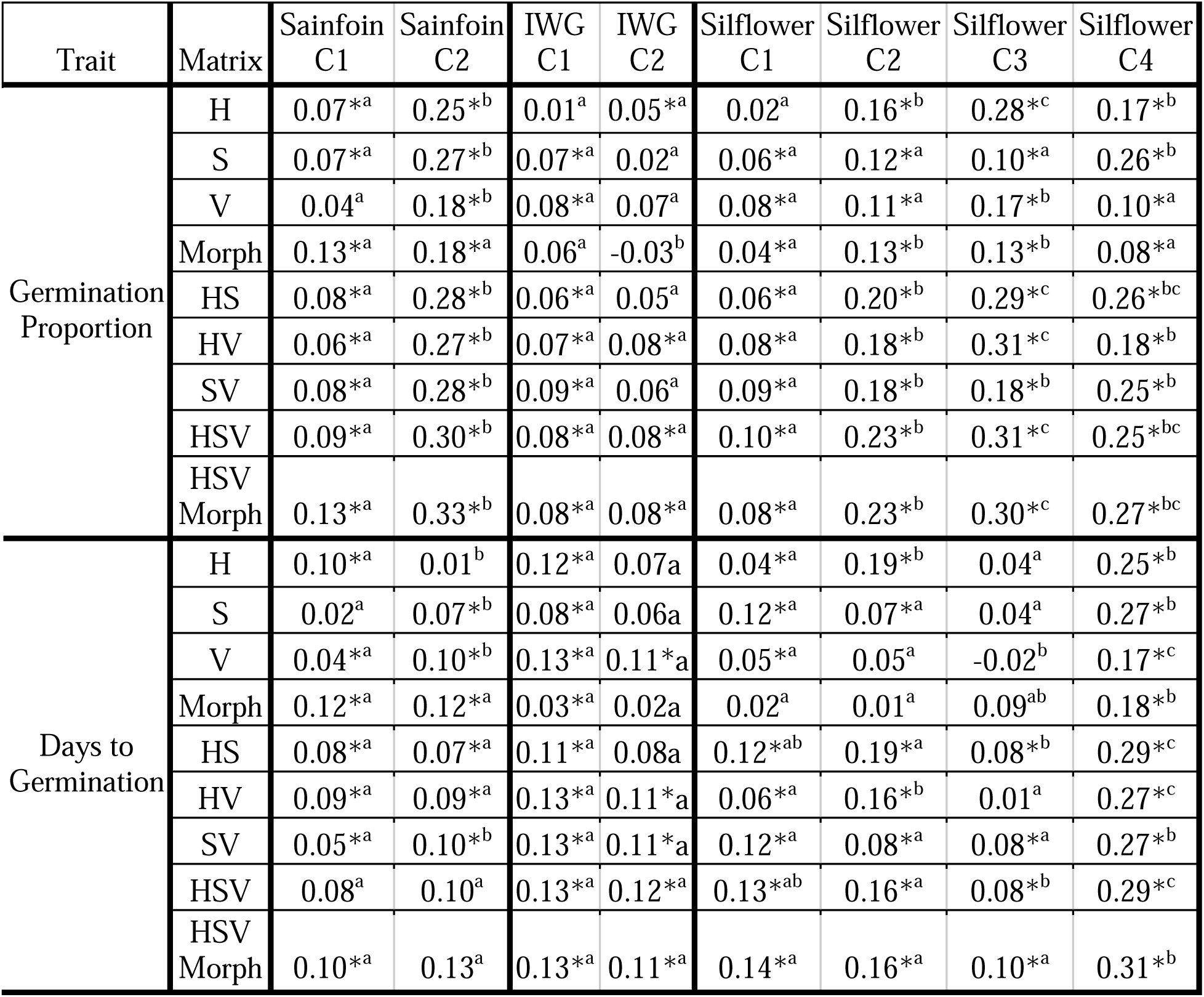
Phenomic selection performance by species and cohort Pearson’s correlation coefficient between observed germination traits and means of the validation set for *Thinopyrum intermedium*, *Onobrychis viciifolia*, and *Silphium integrifolium*. Represents the predictive ability of each relationship matrix. * denotes that the value is significantly different from 0 using a t-test. Letters denote that the cohorts differ significantly using a t-test.

| Trait | Matrix | Sainfoin<br>C1 | Sainfoin<br>C2 | IWG<br>C1 | IWG<br>C2 | Silflower<br>C1 | Silflower<br>C2 | Silflower<br>C3 | Silflower<br>C4 |
| --- | --- | --- | --- | --- | --- | --- | --- | --- | --- |
| Germination<br>Proportion | H | 0.07*<sup>a</sup> | 0.25*<sup>b</sup> | 0.01<sup>a</sup> | 0.05*<sup>a</sup> | 0.02<sup>a</sup> | 0.16*<sup>b</sup> | 0.28*<sup>c</sup> | 0.17*<sup>b</sup> |
|  | S | 0.07*<sup>a</sup> | 0.27*<sup>b</sup> | 0.07*<sup>a</sup> | 0.02<sup>a</sup> | 0.06*<sup>a</sup> | 0.12*<sup>a</sup> | 0.10*<sup>a</sup> | 0.26*<sup>b</sup> |
|  | V | 0.04<sup>a</sup> | 0.18*<sup>b</sup> | 0.08*<sup>a</sup> | 0.07<sup>a</sup> | 0.08*<sup>a</sup> | 0.11*<sup>a</sup> | 0.17*<sup>b</sup> | 0.10*<sup>a</sup> |
|  | Morph | 0.13*<sup>a</sup> | 0.18*<sup>a</sup> | 0.06<sup>a</sup> | -0.03<sup>b</sup> | 0.04*<sup>a</sup> | 0.13*<sup>b</sup> | 0.13*<sup>b</sup> | 0.08*<sup>a</sup> |
|  | HS | 0.08*<sup>a</sup> | 0.28*<sup>b</sup> | 0.06*<sup>a</sup> | 0.05<sup>a</sup> | 0.06*<sup>a</sup> | 0.20*<sup>b</sup> | 0.29*<sup>c</sup> | 0.26*<sup>bc</sup> |
|  | HV | 0.06*<sup>a</sup> | 0.27*<sup>b</sup> | 0.07*<sup>a</sup> | 0.08*<sup>a</sup> | 0.08*<sup>a</sup> | 0.18*<sup>b</sup> | 0.31*<sup>c</sup> | 0.18*<sup>b</sup> |
|  | SV | 0.08*<sup>a</sup> | 0.28*<sup>b</sup> | 0.09*<sup>a</sup> | 0.06<sup>a</sup> | 0.09*<sup>a</sup> | 0.18*<sup>b</sup> | 0.18*<sup>b</sup> | 0.25*<sup>b</sup> |
|  | HSV | 0.09*<sup>a</sup> | 0.30*<sup>b</sup> | 0.08*<sup>a</sup> | 0.08*<sup>a</sup> | 0.10*<sup>a</sup> | 0.23*<sup>b</sup> | 0.31*<sup>c</sup> | 0.25*<sup>bc</sup> |
|  | HSV<br>Morph | 0.13*<sup>a</sup> | 0.33*<sup>b</sup> | 0.08*<sup>a</sup> | 0.08*<sup>a</sup> | 0.08*<sup>a</sup> | 0.23*<sup>b</sup> | 0.30*<sup>c</sup> | 0.27*<sup>bc</sup> |
| Days to<br>Germination | H | 0.10*<sup>a</sup> | 0.01<sup>b</sup> | 0.12*<sup>a</sup> | 0.07<sup>a</sup> | 0.04*<sup>a</sup> | 0.19*<sup>b</sup> | 0.04<sup>a</sup> | 0.25*<sup>b</sup> |
|  | S | 0.02<sup>a</sup> | 0.07*<sup>b</sup> | 0.08*<sup>a</sup> | 0.06<sup>a</sup> | 0.12*<sup>a</sup> | 0.07*<sup>a</sup> | 0.04<sup>a</sup> | 0.27*<sup>b</sup> |
|  | V | 0.04*<sup>a</sup> | 0.10*<sup>b</sup> | 0.13*<sup>a</sup> | 0.11*<sup>a</sup> | 0.05*<sup>a</sup> | 0.05<sup>a</sup> | -0.02<sup>b</sup> | 0.17*<sup>c</sup> |
|  | Morph | 0.12*<sup>a</sup> | 0.12*<sup>a</sup> | 0.03*<sup>a</sup> | 0.02<sup>a</sup> | 0.02<sup>a</sup> | 0.01<sup>a</sup> | 0.09<sup>ab</sup> | 0.18*<sup>b</sup> |
|  | HS | 0.08*<sup>a</sup> | 0.07*<sup>a</sup> | 0.11*<sup>a</sup> | 0.08<sup>a</sup> | 0.12*<sup>ab</sup> | 0.19*<sup>a</sup> | 0.08*<sup>b</sup> | 0.29*<sup>c</sup> |
|  | HV | 0.09*<sup>a</sup> | 0.09*<sup>a</sup> | 0.13*<sup>a</sup> | 0.11*<sup>a</sup> | 0.06*<sup>a</sup> | 0.16*<sup>b</sup> | 0.01<sup>a</sup> | 0.27*<sup>c</sup> |
|  | SV | 0.05*<sup>a</sup> | 0.10*<sup>b</sup> | 0.13*<sup>a</sup> | 0.11*<sup>a</sup> | 0.12*<sup>a</sup> | 0.08*<sup>a</sup> | 0.08*<sup>a</sup> | 0.27*<sup>b</sup> |
|  | HSV | 0.08<sup>a</sup> | 0.10<sup>a</sup> | 0.13*<sup>a</sup> | 0.12*<sup>a</sup> | 0.13*<sup>ab</sup> | 0.16*<sup>a</sup> | 0.08*<sup>b</sup> | 0.29*<sup>c</sup> |
|  | HSV<br>Morph | 0.10*<sup>a</sup> | 0.13<sup>a</sup> | 0.13*<sup>a</sup> | 0.11*<sup>a</sup> | 0.14*<sup>a</sup> | 0.16*<sup>a</sup> | 0.10*<sup>a</sup> | 0.31*<sup>b</sup> |

**Supplemental Table 3:** Phenomic selection performance by matrix

| Trait | contrast | Sainfoin<br>C1 | Sainfoin<br>C2 | IWG<br>C1 | IWG<br>C2 | Silflower<br>C1 | Silflower<br>C2 | Silflower<br>C3 | Silflower<br>C4 |
| --- | --- | --- | --- | --- | --- | --- | --- | --- | --- |
| Germ_prop | H - S | -0.06 | -0.81 | -6.11<br>**** | 1.22 | -1.64 | 1.32 | 6.53 **** | -3.15 * |
| Germ_prop | H - V | 0.98 | 2.54 | -4.07<br>** | -0.64 | -2.37 | 1.85 | 3.96 ** | 2.6 |
| Germ_prop | H - HS | -0.5 | -1.27 | -5.76<br>**** | -0.1 | -1.54 | -1.34 | -0.49 | -3.27 * |
| Germ_prop | H - HV | 0.17 | -0.93 | -3.11 | -1.17 | -2.47 | -0.76 | -0.9 | -0.11 |
| Germ_prop | H - SV | -0.38 | -1.35 | -6.51<br>**** | -0.45 | -2.66 | -0.8 | 3.75 ** | -2.87 |
| Germ_prop | H - HSV | -0.75 | -1.98 | -6.39<br>**** | -0.96 | -2.9 | -2.34 | -1.01 | -2.95 |
| Germ_prop | H - Morph | -2.08 | 2.3 | 5.16<br>**** | 2.82 | -1.03 | 1.07 | 5.32 **** | 3.31 * |
| Germ_prop | H -<br>HSV Morph | -2.33 | -2.97 | -6.58<br>**** | -0.93 | -2.4 | -2.47 | -0.58 | -3.46 * |
| Germ_prop | S - V | 1.04 | 3.35 * | 2.04 | -1.86 | -0.73 | 0.53 | -2.57 | 5.75 **** |
| Germ_prop | S - HS | -0.44 | -0.46 | 0.35 | -1.32 | 0.1 | -2.67 | -7.02<br>**** | -0.12 |
| Germ_prop | S - HV | 0.23 | -0.12 | 3 | -2.39 | -0.83 | -2.09 | -7.43<br>**** | 3.04 |
| Germ_prop | S - SV | -0.32 | -0.54 | -0.39 | -1.67 | -1.02 | -2.12 | -2.78 | 0.28 |
| Germ_prop | S - HSV | -0.69 | -1.17 | -0.28 | -2.18 | -1.26 | -3.67 ** | -7.55<br>**** | 0.2 |
| Germ_prop | S - Morph | -2.02 | 3.1 | 11.27<br>**** | 1.6 | 0.61 | -0.26 | -1.21 | 6.46 **** |
| Germ_prop | S -<br>HSV Morph | -2.28 | -2.16 | -0.46 | -2.15 | -0.76 | -3.8 ** | -7.12<br>**** | -0.32 |
| Germ_prop | V - HS | -1.48 | -3.8 ** | -1.69 | 0.54 | 0.83 | -3.19 * | -4.45 *** | -5.87<br>**** |
| Germ_prop | V - HV | -0.81 | -3.47 * | 0.96 | -0.53 | -0.1 | -2.61 | -4.86<br>**** | -2.71 |
| Germ_prop | V - SV | -1.36 | -3.88 ** | -2.44 | 0.19 | -0.29 | -2.65 | -0.21 | -5.46<br>**** |
| Germ_prop | V - HSV | -1.73 | -4.52 *** | -2.32 | -0.32 | -0.54 | -4.19 ** | -4.97<br>**** | -5.55<br>**** |
| Germ_prop | V - Morph | -3.06 | -0.24 | 9.23<br>**** | 3.46 * | 1.34 | -0.78 | 1.36 | 0.71 |
| Germ_prop | V -<br>HSV Morph | -3.32 * | -5.5 **** | -2.51 | -0.29 | -0.04 | -4.32 *** | -4.54 *** | -6.06<br>**** |
| Germ_prop | HS - HV | 0.67 | 0.34 | 2.65 | -1.08 | -0.93 | 0.58 | -0.41 | 3.16 * |
| Germ_prop | HS - SV | 0.12 | -0.08 | -0.75 | -0.36 | -1.12 | 0.55 | 4.24 *** | 0.41 |
| Germ_prop | HS - HSV | -0.25 | -0.71 | -0.63 | -0.87 | -1.37 | -1 | -0.53 | 0.32 |
| Germ_prop | HS - Morph | -1.58 | 3.56 * | 10.92 **** | 2.92 | 0.51 | 2.41 | 5.81 **** | 6.58 **** |
| Germ_prop | HS - HSV Morph | -1.84 | -1.7 | -0.82 | -0.83 | -0.87 | -1.13 | -0.1 | -0.19 |
| Germ_prop | HV - SV | -0.55 | -0.42 | -3.4 * | 0.72 | -0.19 | -0.03 | 4.65 *** | -2.75 |
| Germ_prop | HV - HSV | -0.92 | -1.05 | -3.28 * | 0.21 | -0.43 | -1.58 | -0.12 | -2.84 |
| Germ_prop | HV - Morph | -2.25 | 3.22 * | 8.27 **** | 3.99 ** | 1.45 | 1.83 | 6.22 **** | 3.42 * |
| Germ_prop | HV - HSV Morph | -2.51 | -2.04 | -3.47 * | 0.24 | 0.07 | -1.71 | 0.31 | -3.35 * |
| Germ_prop | SV - HSV | -0.37 | -0.63 | 0.12 | -0.51 | -0.24 | -1.55 | -4.77 **** | -0.08 |
| Germ_prop | SV - Morph | -1.7 | 3.64 ** | 11.67 **** | 3.27 * | 1.64 | 1.86 | 1.57 | 6.18 **** |
| Germ_prop | SV - HSV Morph | -1.96 | -1.62 | -0.07 | -0.48 | 0.26 | -1.68 | -4.34 *** | -0.6 |
| Germ_prop | HSV - Morph | -1.33 | 4.28 *** | 11.55 **** | 3.78 ** | 1.88 | 3.41 * | 6.34 **** | 6.26 **** |
| Germ_prop | HSV - HSV Morph | -1.58 | -0.99 | -0.19 | 0.03 | 0.5 | -0.13 | 0.43 | -0.51 |
| Germ_prop | Morph - HSV Morph | -0.25 | -5.26 **** | -11.74 **** | -3.75 ** | -1.38 | -3.54 * | -5.9 **** | -6.77 **** |
| Days_to_germ | H - S | 2.63 | -2.08 | 0.74 | 0.11 | -2.75 | 4.27 *** | -1.67 | -0.52 |
| Days_to_germ | H - V | 1.98 | -3.33 * | 0.03 | -1.48 | -0.12 | 4.91 **** | 2.17 | 2.88 |
| Days_to_germ | H - HS | 0.47 | -2.35 | 0.09 | -0.42 | -2.85 | 0.05 | -1.29 | -1.25 |
| Days_to_germ | H - HV | 0.32 | -2.9 | 0.02 | -1.59 | -0.69 | 1.07 | 1.03 | -0.73 |
| Days_to_germ | H - SV | 1.85 | -3.51 * | 0.06 | -1.52 | -2.83 | 4.01 ** | -1.58 | -0.38 |
| Days_to_germ | H - HSV | 0.53 | -3.36 * | 0.03 | -1.73 | -2.96 | 0.91 | -1.36 | -1.41 |
| Days_to_germ | H - Morph | -0.67 | -4 ** | 2.37 | 1.56 | 0.93 | 6.57 **** | -1.72 | 2.83 |
| Days_to_germ | H - HSV Morph | -0.28 | -4.3 *** | -0.12 | -1.62 | -3.35 * | 1.14 | -2.05 | -1.96 |
| Days_to_germ | S - V | -0.64 | -1.25 | -0.71 | -1.59 | 2.62 | 0.64 | 3.85 ** | 3.4 * |
| Days_to_germ | S - HS | -2.16 | -0.27 | -0.65 | -0.53 | -0.1 | -4.22 *** | 0.38 | -0.74 |
| Days_to_germ | S - HV | -2.31 | -0.81 | -0.71 | -1.7 | 2.05 | -3.2 * | 2.7 | -0.21 |
| Days_to_germ | S - SV | -0.78 | -1.43 | -0.68 | -1.62 | -0.08 | -0.26 | 0.09 | 0.13 |
| Days_to_germ | S - HSV | -2.09 | -1.28 | -0.7 | -1.84 | -0.21 | -3.36 * | 0.31 | -0.9 |
| Days_to_germ | S - Morph | -3.3 * | -1.92 | 1.63 | 1.46 | 3.68 ** | 2.3 | -0.05 | 3.35 * |
| Days_to_germ | S - HSV Morph | -2.91 | -2.22 | -0.86 | -1.72 | -0.6 | -3.13 * | -0.37 | -1.44 |
| Days_to_germ | V - HS | -1.51 | 0.98 | 0.06 | 1.07 | -2.72 | -4.86 **** | -3.46 * | -4.14 ** |
| Days_to_germ | V - HV | -1.67 | 0.43 | -0.01 | -0.1 | -0.57 | -3.84 ** | -1.14 | -3.62 ** |
| Days_to_germ | V - SV | -0.13 | -0.18 | 0.03 | -0.03 | -2.7 | -0.9 | -3.76 ** | -3.27 * |
| Days_to_germ | V - HSV | -1.45 | -0.04 | 0 | -0.24 | -2.83 | -4 ** | -3.54 * | -4.3 *** |
| Days_to_germ | V - Morph | -2.66 | -0.68 | 2.34 | 3.05 | 1.06 | 1.66 | -3.89 ** | -0.05 |
| Days_to_germ | V - HSV Morph | -2.26 | -0.97 | -0.15 | -0.13 | -3.23 * | -3.77 ** | -4.22 *** | -4.84 **** |
| Days_to_germ | HS - HV | -0.15 | -0.54 | -0.06 | -1.17 | 2.15 | 1.02 | 2.32 | 0.52 |
| Days_to_germ | HS - SV | 1.38 | -1.15 | -0.03 | -1.1 | 0.02 | 3.96 ** | -0.29 | 0.87 |
| Days_to_germ | HS - HSV | 0.06 | -1.01 | -0.05 | -1.31 | -0.11 | 0.86 | -0.07 | -0.16 |
| Days_to_germ | HS - Morph | -1.14 | -1.65 | 2.28 | 1.98 | 3.78 ** | 6.52 **** | -0.43 | 4.08 ** |
| Days_to_germ | HS - HSV Morph | -0.75 | -1.95 | -0.2 | -1.2 | -0.5 | 1.09 | -0.76 | -0.7 |
| Days_to_germ | HV - SV | 1.53 | -0.61 | 0.04 | 0.07 | -2.13 | 2.94 | -2.61 | 0.35 |
| Days_to_germ | HV - HSV | 0.22 | -0.47 | 0.01 | -0.14 | -2.26 | -0.17 | -2.39 | -0.68 |
| Days_to_germ | HV - Morph | -0.99 | -1.11 | 2.35 | 3.15 * | 1.63 | 5.5 **** | -2.75 | 3.56 * |
| Days_to_germ | HV - HSV Morph | -0.6 | -1.41 | -0.14 | -0.03 | -2.66 | 0.07 | -3.08 | -1.23 |
| Days_to_germ | SV - HSV | -1.32 | 0.14 | -0.03 | -0.21 | -0.13 | -3.11 | 0.22 | -1.03 |
| Days_to_germ | SV - Morph | -2.52 | -0.5 | 2.31 | 3.08 | 3.76 ** | 2.56 | -0.14 | 3.21 * |
| Days_to_germ | SV - HSV Morph | -2.13 | -0.8 | -0.18 | -0.1 | -0.52 | -2.87 | -0.46 | -1.57 |
| Days_to_germ | HSV - Morph | -1.2 | -0.64 | 2.34 | 3.29 * | 3.89 ** | 5.66 **** | -0.36 | 4.24 *** |
| Days_to_germ | HSV - HSV Morph | -0.81 | -0.94 | -0.15 | 0.11 | -0.39 | 0.23 | -0.68 | -0.54 |
| Days_to_germ | Morph - HSV Morph | 0.39 | -0.3 | -2.49 | -3.18 * | -4.28 *** | -5.43 **** | -0.33 | -4.79 **** |

